# Tau spreading coordinates intercellular lipid flux enhancing neuronal resilience to lipid toxicity

**DOI:** 10.64898/2026.07.30.741734

**Authors:** Anna Oliveras Martinez, Davide Ibatici, Maria Calvo Noriega, Guido Mastrobuoni, Martin Forbes, Leandro Manuel Santiago Padilla, Melisa Aykul, Kamelija Horvatovic, Mara-Camelia Rusu, Melina Patsonis, Laurent Jutras-Dubé, Jana Rossius, Cledi Cerda Jara, Ella Bahry, Johannes Broichhagen, Deborah Schmidt, Jakob Metzger, Nikolaus Rajewsky, Séverine Kunz, Agnieszka Rybak-Wolf, Stefan Kempa, Melissa Birol

**Affiliations:** Laboratory for Single Molecule Biophysics probing Quantitative Neuroscience, Berlin Institute for Medical Systems Biology, Max Delbrück Center for Molecular Medicine in the Helmholtz Association, Berlin, Germany; Laboratory for Integrative Proteomics and Metabolomics, Berlin Institute for Medical Systems Biology, Max Delbrück Center for Molecular Medicine in the Helmholtz Association, Berlin, Germany; Berlin Institute of Health at Charité – Universitätsmedizin Berlin, Berlin, Germany; Electron Microscopy Technology Platform, Max Delbrück Center for Molecular Medicine in the Helmholtz Association, Berlin, Germany; Laboratory for Quantitative Stem Cell Biology, Berlin Institute for Medical Systems Biology, Max Delbrück Center for Molecular Medicine in the Helmholtz Association, Berlin, Germany; Laboratory for Systems Biology of Gene Regulatory Elements, Berlin Institute for Systems Biology, Max Delbrück Center for Molecular Medicine, Berlin, Germany; Helmholtz Imaging, Max Delbrück Center for Molecular Medicine in the Helmholtz Association, Berlin, Germany; Leibniz-Forschungsinstitut für Molekulare Pharmakologie (FMP), Berlin, Germany; Charité – Universitätsmedizin, Berlin, Germany; German Center for Cardiovascular Research (DZHK), Berlin, Germany; German Cancer Consortium (DKTK), Heidelberg, Germany; NeuroCure Cluster of Excellence, Berlin, Germany; National Center for Tumor Diseases (NCT), Berlin, Germany; ImmunoPreCept, Cluster of Excellence, Berlin, Germany; Einstein Center for Early Disease Interception, Berlin, Germany; Organoid Platform, Berlin Institute for Medical Systems Biology, Max Delbrück Center for Molecular Medicine in the Helmholtz Association, Berlin, Germany

**Author notes:** Equal contribution (Davide Ibatici, Maria Calvo Noriega). Address correspondence to: Melissa Birol.

## Abstract

The presence of intrinsically disordered proteins (IDPs) in the extracellular environment of the brain suggests that protein disorder may serve functions beyond those confined to the intracellular space. Tau, a prototypical IDP linked to tauopathies, a class of neurodegenerative diseases, is continuously released and spreads between cells in the brain, yet the biological significance of this extracellular phase remains unresolved. Here, we show that spreading tau coordinates neutral lipid homeostasis by promoting neuronal lipid efflux and enhancing resilience to lipid toxicity. Integrating transcriptomics and lipidomics, we demonstrate that tau spreading reprograms lipid transport pathways and triggers redistribution of triacylglycerol pools, remodeling neutral lipid metabolism. Mechanistically, cellular uptake of spreading tau drives its accumulation within neutral lipid-rich compartments and promotes neuronal lipid efflux. The exported lipids are enriched in peroxidized species that are subsequently transferred to astrocytes, reducing neuronal lipid stress and revealing a pathway through which tau regulates lipid homeostasis. Our findings reframe spreading tau as a regulator of intercellular lipid flux and position protein disorder as an active mediator of cell-to-cell communication in coordinating tissue metabolism. Loss of this homeostatic function is likely to contribute to the early metabolic dysfunction associated with tauopathy.

## Introduction

Intrinsically disordered proteins (IDPs) lack a tertiary structure yet execute diverse functions through multivalent, transient interactions^1–5^. Although historically viewed as intracellular regulators, many IDPs are secreted and persist in the extracellular space. Secreted IDPs are released from the brain^6,7^, liver^8,9^ pancreas^10^, immune tissues^11^, and endocrine glands^12^ via both classical ER-Golgi trafficking and unconventional pathways such as exosomes, ectodomain shedding, and stress-induced leakage^13–15^. Many of these peripheral IDPs act as hormone-like modulators, antimicrobial factors, or matricellular scaffolds that exploit conformational plasticity to engage a broad range of extracellular partners^1^.Dysregulated secretion or misfolding of IDPs contributes to diverse pathologies, including metabolic dysfunction, systemic amyloidosis, and chronic inflammatory states^16–20^. In the brain, they are found in the interstitial fluid, cerebrospinal fluid (CSF), presynaptic milieu, and extracellular matrix (ECM)^21–28^. In these spaces, they are proposed to modulate synaptic vesicle release, neuronal synaptic signaling and extracellular organization through multivalent interactions and condensate formation^20,29,30^. However, despite their widespread extracellular presence, the biological significance of this extracellular phase, in particular prior to aggregation, remains poorly defined. This knowledge gap is especially relevant in neurodegeneration, where extracellular protein spreading is link to disease progression. Determining whether extracellular IDPs serve physiological functions is therefore essential for understanding how normal protein signaling transitions into pathological propagation.

Tau represents a compelling model for investigating the extracellular functions of IDPs in the brain^31^. Encoded by MAPT, tau exists as six splice isoforms that differ in N-terminal inserts (N) and microtubule-binding repeats (R)^32^. All isoforms are expressed in the adult human brain, where the predominant populations are the 3-R repeat (2N3R) and 4-R repeat (2N4R) isoforms, present at an approximately 1:1 ratio^33^. Beyond its canonical intracellular roles in microtubule dynamics and axonal transport^34,35^, tau undergoes intercellular transfer through a process we refer to as tau spreading, encompassing its release from donor cells, transit through the extracellular space, and uptake by recipient cells. Consistent with this, tau is continuously released under physiological conditions and is detectable in the interstitial fluid and CSF of healthy individuals ^7,28,33,36–38^. Notably, extracellular tau levels are dynamically regulated and exhibit disease- and compartment-specific alterations across tauopathies^39^. In tauopathies such as Alzheimer’s disease (AD) and frontotemporal lobar degeneration (FTLD), tau undergoes aberrant post-translational modifications, dissociates from microtubules, and assembles into oligomers and fibrils that propagate across synaptically connected regions^40,41^. Notably, distinct tauopathies exhibit characteristic spatiotemporal patterns of tau spread through the brain^42,43^, suggesting that propagation follows defined biological pathways rather than occurring randomly. This process involves extracellular trafficking of soluble and aggregated tau species via defined receptor-mediated pathways, including heparan sulfate proteoglycans (HSPGs) and LRP1 receptor^44–46^. While tau propagation, specifically in its aggregated form, has been extensively studied in the context of neurodegeneration and disease progression, the functional significance of its extracellular phase prior to pathology remains unclear.

Neutral lipid metabolism represents a major axis of vulnerability in neurodegeneration^47,48^. Neutral lipids, primarily triacylglycerols (TGs) and cholesteryl esters (CEs), are stored in lipid droplets (LDs)^49^, and their homeostasis depends on dynamic neuron-astrocyte metabolic coupling. Under elevated activity, neurons generate reactive oxygen species (ROS) that peroxidate polyunsaturated fatty acids (PUFAs), yielding free fatty acids (FAs) and lipid peroxides. To compensate for their limited storage and catabolic capacity, neurons export toxic lipids to astrocytes via apolipoprotein E (APOE)-containing lipoprotein particles^50,51^. APOE is the major lipid carrier in the brain and APO4 isoform the strongest genetic risk factor for sporadic Alzheimer’s disease ^52,53^. Disruption of this intercellular lipid flux, prominent in APOE4 carriers, compromises astrocytic buffering capacity and is associated with elevated lipid peroxidation in AD brain and CSF ^54–56^. Neutral lipids accumulate in early tauopathy and neuroinflammation ^57–60^ and markers of lipid peroxidation are elevated in AD brain and CSF ^61,62^. In human AD tissue and models, neutral lipids and LDs are enriched around plaques and in glial cells, indicating broad disturbances in neutral lipid storage, turnover or neuron-astrocyte coupling ^59,63–65^. Accumulating evidence further links tau pathology to disruption of brain lipid homeostasis ^59,66–68^. Tau pathology is sufficient to induce lipid dysregulations even in the absence of amyloid deposition ^67,69,70^ and disease progression is accompanied by early changes in lipid composition and LD dynamics^66^. Lipid disturbances influence tau aggregation, seeding, and propagation, highlighting a bidirectional relationship between tau and lipid homeostasis ^66^. Moreover, APOE4 exacerbates tau pathology ^71–74^ while LD-rich microglial states promote tau phosphorylation and neurotoxicity in an APOE-dependent manner ^59^. Glial tau is also required for LD formation and protection against neuron-derived peroxidized lipids ^75^. These findings suggest a mechanistic intersection between tau biology and lipid homeostasis. However, the physiological significance of tau spreading and its potential impact on brain metabolism remain poorly understood.

In this study, we uncover a previously unrecognized function of spreading tau as a regulator of neuronal lipid metabolism, protecting against lipid toxicity. Through integrated lipidomic and transcriptomic analyses of tau-recipient and non-recipient cells, we show that tau spreading triggers selective remodeling of neutral lipid pools and coordinated changes of lipid regulatory pathways. Mechanistically, spreading tau localizes to intracellular neutral lipid, enhances FA efflux, and facilitates transfer of toxic lipid species from neurons to astrocytes. This tau-driven lipid redistribution reinforces intercellular lipid coupling across brain tissue and reduces oxidative burden. Our findings establish extracellular tau transmission as a signaling process that reshapes brain lipid metabolism and intercellular crosstalk. The dysregulation of this homeostatic pathway may contribute to early metabolic dysfunction in tauopathy, linking protein spreading to the maintenance of metabolic balance in the brain.

## Results

### Progressive cell-to-cell spreading of tau

To model the spontaneous spread of monomeric tau within brain organoids, we conditioned forebrain organoids with recombinantly expressed full-length human tau (2N4R). Endogenous cysteines (C291 and C322) were mutated, and a single cysteine was introduced at the C-terminus (residue 433) to enable site-specific labeling with maleimide-conjugated Alexa-647, generating fluorescent tau (tau_A647_). Following validation of functional neuronal activity by calcium imaging (Fig. S1a-c, movie 1), the organoid media was supplemented with 20 nM tau_A647_ monomer protein. This concentration is below estimated endogenous intracellular tau levels in neurons (∼2 μM)^76,77^, yet several orders of magnitude higher than bulk CSF tau concentrations, which lie in the low-picomolar range ^78^ Organoids were exposed to tau_A647_ for 16h to permit cellular uptake, after which non-internalized excess extracellular tau was washed out. This washout point was defined as t = 0, and intercellular tau spreading was monitored over the subsequent 24, 48, and 120 h (Fig. 1a). Calcium imaging showed no significant differences in spontaneous neuronal activity between control and tau-treated organoids, as assessed by calcium firing frequency and peak amplitude (Fig. S1a- e, movie 1). Similarly, tau conditioning did not affect cell viability (Fig. S1f), indicating that tau spreading occurs in the absence of neuronal dysfunction and toxicity.

**Figure 1.**
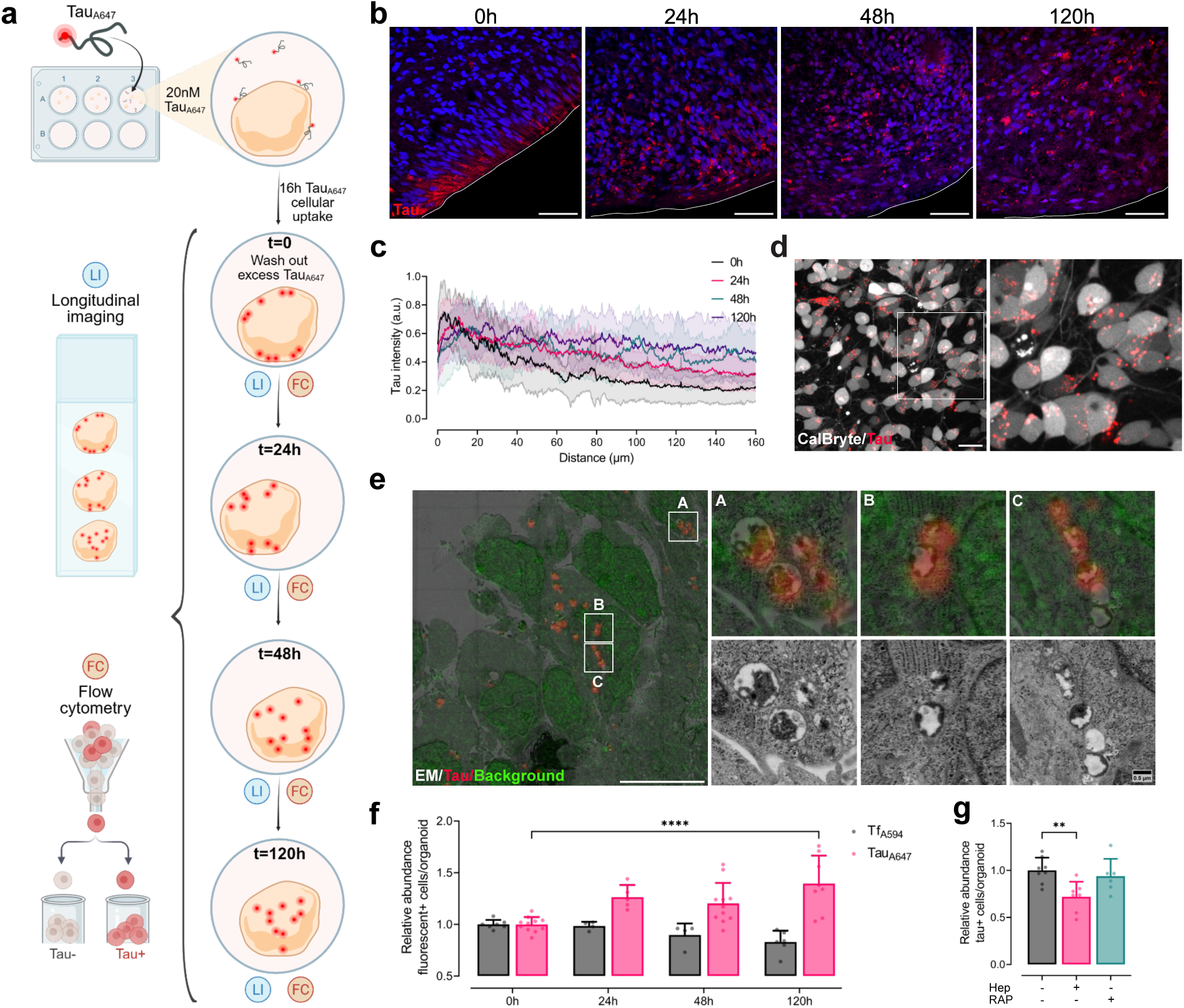
Spatiotemporal spreading dynamics of monomeric tau across cells in human forebrain organoids. (a) Schematic of experimental setup of tau spreading model. (b) Confocal images of organoid sections showing spatial redistribution of tau_A647_ (red) over time (0, 24, 48 and 120h). Nuclei in blue. Scale bars, 50 µm. (c) Quantification of radial tau_A647_ intensity profiles normalized to the maximum intensity value of each line profile at the different time points of spreading in (b). Each line represents the mean ± SD, *n* = 10-17 images, 3 experimental replicates. (d) Left: representative frame of live-imaging time-lapse experiment on tau_A647_–treated organoid. Tau is shown in red and cell bodies can be recognized by CalBryte520 signal (grey-scale). Right: boxed area is shown at higher magnification. Scale bar, 20 µm. (e) Left: Overview correlative fluorescence and EM of organoids 48 h after tau_A647_ exposure: in grey-scale, EM image; in red, fluorescent tau; in green, fluorescent background signal. Boxed areas highlight representative micrometer- sized ROIs. Scale bar, 10 µm. Right: Boxed ROIs shown at higher magnification highlighting tau assemblies. Corresponding correlated and scanning electron microscopy micrographs are shown. Scale bar, 0.5 µm. 3 experimental replicates (f) Relative abundance of tau_A647_+ and transferrin_A594_ (Tf_A594_+) cells per organoid at increasing time points. Data were normalized to the initial uptake time point, t=0. For tau_A647_, *n* = 5-11 individual organoids per time point, 4 independent experiments. For Tf_A594_ *n* = 4-7 individual organoids per time point, 3 independent experiments. Data were analyzed by two-way ANOVA, interaction *p* = 0.0001 and with protein (*p* < 0.0001) and time point (ns) as independent variables, post-hoc Tukey’s tau_A647_ 0 h versus 120 h, *p* < 0.0001. (g) Percentage of tau_A647_+ cells at 120 h in untreated (-Hep) and heparinase I/III treated organoids (+Hep). For -Hep, *n* = 6 individual organoids and for +Hep, *n* = 4, 2 experimental replicates. P value was determined by Welch’s test.

We next examined the spatial dynamics of tau spreading within intact organoids. Longitudinal imaging of organoid slices revealed a progressive inward translocation of tau_A647_ from the outer cellular layers, where tau uptake occurs at t=0, toward the organoid core over subsequent time points (Fig. 1b). This redistribution was accompanied by a gradual expansion of tau_A647_ intensity from the periphery to the inner layers. By 120h, tau signal was distributed throughout both peripheral and inner regions extending to a depth of approximately 215 ± 19 µm, corresponding to 12 ± 4 cell layers from the surface (Fig. 1b, c). Immunostaining for cell-type-specific markers showed tau_A647_ positive-progenitors (SOX2⁺), neurons (βIII-tubulin⁺), and astrocytes (S100B⁺), detecting spreading tau in multiple neural cell populations within the organoid (Fig. S1g). Time-lapse imaging of live tau_A647_-treated organoids further revealed that intracellular tau_A647_ assemblies are dynamic and undergo directed motion within cell bodies, with an average velocity of 0.24 µm/s (Fig. 1d, movie 2). To define the ultrastructure of these tau assemblies, we performed correlative light and electron microscopy (CLEM), combining confocal fluorescence imaging with scanning electron microscopy (Fig. 1e, S1h). Spreading tau accumulated in discrete intracellular puncta distributed across cell bodies and neurites, with an average cross-sectional area of 0.29 ± 0.21 µm² (n=35). Ultrastructural analysis revealed that these puncta correspond to membrane-bound compartments with heterogeneous internal organization, characterized by regions of electron-dense and electron-lucent material. Electron-dense material was predominantly enriched near the limiting membrane, with occasional extensions into the interior of the compartment. Despite progressive intracellular accumulation of tau puncta, no Thioflavin S (ThioS)-positive aggregates were detected in organoids conditioned with tau_A594_ compared to organoids treated with tau fibers_A594_ (Fig. S1i), and tau phosphorylation levels, assessed by AT8 immunostaining, remained comparable to control untreated organoids (Fig. S1j-k). These data support that internalized tau forms dynamic, membrane-associated assemblies while remaining non-fibrillar.

To quantify tau propagation, organoids were dissociated into single cells at each time point and subjected to flow cytometry to determine the fraction of tau_A647_-positive cells (Fig. 1f). As a control for uptake and turnover, organoids were treated with Alexa594-labeled transferrin (Tf_A594_). While the proportions of tau_A647_ and Tf_A594_-positive cells at t=0, were comparable (54 ± 7% and 67 ± 17%, respectively), the relative abundance of Tf_A594_-positive cells declined over time, consistent with transferrin degradation. In contrast, the relative abundance of tau_A647_-positive cells steadily increased over 120 h, reaching significantly higher levels at 120 h compared with t=0 (t=0: 1.0 ± 0.1 a.u. vs. t=120h: 1.4 ± 0.3 a.u., *p* < 0.0001), consistent with active intercellular spreading. The 24h and 48h (60 ± 5 % and 63 ± 8% of tau_A647_-positive cells, respectively) time points were selected for most subsequent experiments unless otherwise specified. To test whether established mediators of tau uptake contribute to intercellular spreading, we enzymatically depleted cell-surface heparan sulfate proteoglycans (HSPG) using heparinases I/III, and inhibited LRP1 using the receptor-associated protein (RAP), following the time of initial tau exposure (t = 0, Fig. 1g). When assessed at 120 h and normalized to the corresponding untreated (-Hep/-RAP) condition, HSPG depletion markedly suppressed tau spreading, limiting the relative abundance of tau_A647_-positive cells (-Hep/-RAP: 1.0± 0.1 a.u. vs. + Hep: 0.7 ± 0.2 a.u., *p* = 0.0081), whereas RAP treatment did not significantly perturb tau spreading (+RAP: 0.9 ± 0.2 a.u., p = 0.7600). Thus, HSPGs, but not LRP1, are required not only for tau uptake but for its progressive cell-to-cell dissemination in tissue. Together, these data establish a human brain organoid system that permits controlled analysis of progressive spreading of initially monomeric tau and its assembly into early assemblies prior to aggregation.

### Tau spreading reprograms lipid metabolism gene networks

Having established that tau spreads as dynamic, non-fibrillar assemblies across multiple neural cell types, we next asked whether this process alters cellular metabolic state within tau spreading trajectories. We performed RNA sequencing on cell populations from untreated organoids and tau_A647_-conditioned organoids, FACS-sorted into tau-containing cells (tau_A647_+) and tau-exposed but non-internalizing cells (tau_A647_-). Principal component analysis (PCA) revealed clear segregation of all three populations (Fig. 2a), suggesting that tau spreading induces strong and system-wide transcriptional changes in both tau_A647_+ (3,017 DEGs compared to control) and in tau_A647_- (15,904 DEGs compared to control). We first examined transcriptional changes at the whole-organoid level (Fig. S2a). Consistent with previously described functions of tau, gene sets associated with microtubule organization, chromatin regulation, and cellular stress responses were altered in tau-conditioned organoids ^79–82^ Notably, metabolic pathways, including FA metabolism, eicosanoid metabolism, and superoxide metabolic processes were also enriched, indicating engagement of lipid-related oxidative stress programs at the tissue level. Enrichment of inflammatory responses pathways further indicate engagement of immune-related signaling.

**Figure 2.**
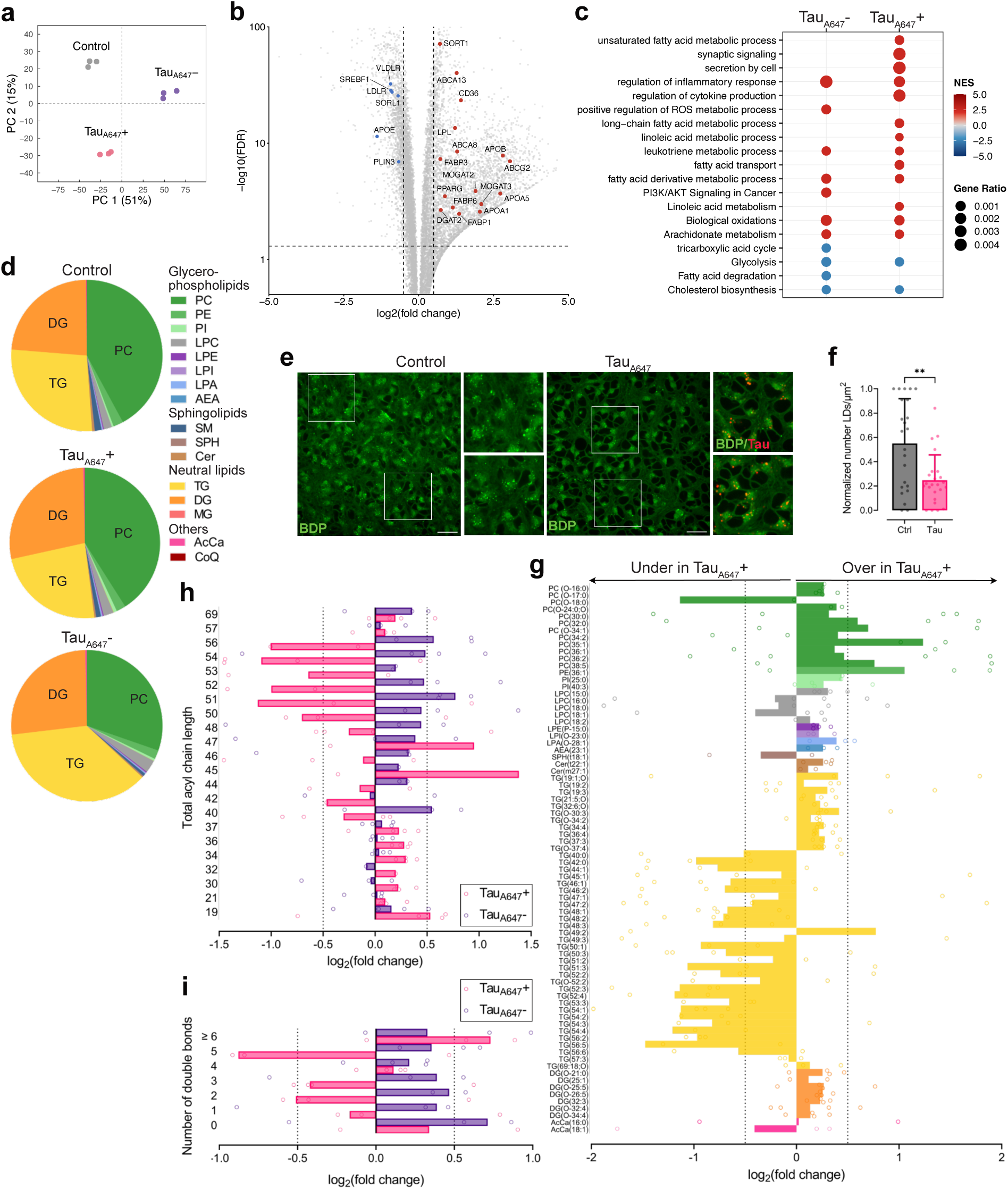
Tau spreading establishes distinct metabolic states and remodels neutral lipid metabolism. (a) PCA analysis of transcriptomic data comparing control, tau_A647_+ and tau_A647_- cell populations. (b) Volcano plot of genes differentially expressed in tau_A647_-conditioned organoid relative to control (log₂ fold change versus -log₁₀ adjusted p value). Relevant lipid metabolism-associated genes are highlighted in red (upregulated) and blue (downregulated). Dashed lines denote statistical and fold-change cutoffs. (c) GSEA dot plot showing selected significantly enriched pathways in tau_A647_+ and tau_A647_- cells compared to control. Dot size represents gene ratio; color indicates normalized enrichment score (NES). For all transcriptomics data (a-c), data represents *n* = 3 technical replicates. (d) Pie charts showing the distribution of major lipid classes in control, tau_A647_+ and tau_A647_- cell populations (e) Live-cell imaging of control and tau-conditioned organoids with tau_A647_ (red) and co-stained with BDP (green). White squares highlight the areas shown at higher magnification on the right. Scale bars, 20 µm (f) Quantification of the number of LDs/μm^2^ in (e). Values are shown as mean after min-max normalization across independent experiments. *n* = 23 (Ctrl) and *n* = 24 (Tau) FOVs, 5 independent experiments. P value (*p* = 0.0089) was determined by a Mann-Whitney U test. (g) Lipid species-resolved log₂ fold change (tau_A647_+ versus tau_A647_-) of lipid intensities for all detected lipids, ordered by class and color-coded, sub-ordered by acyl chain length. Bars represent the mean log₂ fold changes for each experimental replicate (dots). (h) Log₂ fold change of TG species grouped by total acyl chain length, comparing tau_A647_+ cells and tau_A647_- cells to control. Dots represent the sum of lipid intensities of all TG species with given acyl chain length for each experimental replicate (i) Log₂ fold change of TGs stratified by degree of saturation, comparing tau_A647_+ cells and tau_A647_- cells to control. Dots represent the sum of lipid intensities of all TG species for a given degree of saturation for each experimental replicate. For all lipidomics, (d), (g-i), data represent independent organoid batches, *n* = 4 experimental replicates.

Differential expression analysis revealed a transcriptional program consistent with active lipid mobilization (Fig. 2b). Genes mediating lipid transport and efflux including, SORT1, ABCG2, ABCA13 and multiple apolipoproteins (APOA1, APOA4, APOA5), were upregulated. In parallel, genes involved in lipid uptake and intracellular handling, such as CD36, LPL, FABP4/5, as well as genes involved in FA esterification, including MOGAT2/3 and DGAT2 were also induced. In contrast, genes associated with lipid synthesis and storage were downregulated, including SREBF1, a master regulator of lipogenesis, and PLIN3, the predominant perilipin coating neuronal LDs. APOE, and several of its canonical receptors (SORL1, LDLR and VLDLR) were likewise downregulated. This transcriptional program reflects activation of FA mobilization and export pathways despite reduced expression of APOE and its canonical receptors, suggesting engagement of alternative lipid transport mechanisms.

Direct comparison of tau_A647_+ and tau_A647_- cells revealed striking divergence in metabolic and signaling programs between the two populations (Fig. 2c), indicating that tau spreading partitions neural tissue into distinct functional states. Tau_A647_+ cells were enriched for pathways related to long-chain, unsaturated, and linoleic FA metabolism, FA transport, eicosanoid and leukotriene metabolism, biological oxidation, synaptic signaling, and cellular secretion. These pathways support a role for tau spreading in neutral lipid mobilization, PUFA release, and inflammatory stress-responsive lipid handling. In contrast, tau_A647_- cells showed enrichment of PI3K-AKT signaling and ROS metabolic processes, together with suppression of cholesterol biosynthesis, central carbon metabolism, and tricarboxylic acid cycle, and FA degradation. This transcriptional profile suggests reduced engagement of anabolic metabolic pathways and activation of oxidative stress-adaptive signaling.

### Tau spreading selectively remodels neutral lipid metabolism

To determine whether these transcriptional changes are accompanied by altered lipid composition, we performed untargeted lipidomics on the same populations. Comparison of global lipid class distributions between control, tau_A647_+ and tau_A647_- revealed a pronounced and selective reorganization of neutral lipid metabolism (Fig. 2d and Table1). Relative to control, tau_A647_+ cells showed a marked reduction in total TG abundance, whereas TG levels were substantially elevated in tau_A647_- cells. Among all detected lipid classes, TGs exhibited the most pronounced and consistent fold changes, accounting for ∼20-40% of total lipids and comprising over 40 molecular species. This identifies TGs as the dominant contributors to tau-associated lipid remodeling (Fig. S2b). Levels of phosphatidylcholine (PC), the most abundant phospholipid class, were comparable between tau_A647_+ and control cells. In contrast, tau_A647_- cells displayed a modest reduction in PC, likely reflecting the increased contribution of TGs to the total lipid pool in this population (Fig. 2d and Table 1). Other major phospholipid classes, including phosphatidylethanolamine (PE) and several lysophospholipid derivates, as well as major sphingolipids classes, remined largely unchanged across conditions (Table1). To validate changes in the TG pool at the tissue level, organoids were stained with the neutral lipid dye BODIPY 493/503 and analyzed by spot detection (Fig. 2e, S2c). Tau_A647_+ tissue displayed a reduced abundance of LDs relative to controls (Fig. 2f), supporting that the TG depletion detected by lipidomics is accompanied by reduced neutral lipid storage in tau-internalizing cells.

**Table 1.** Relative abundance of all detected lipid classes.

| Lipid classes | Control | Tau<sub>A647</sub><sup>+</sup> | Tau<sub>A647</sub><sup>-</sup> |
| --- | --- | --- | --- |
| PC | 42.09 ± 18.57 | 40.98 ± 24.75 | 30.31 ± 23.78 |
| PE | 2.15 ± 1.40 | 2.02 ± 2.07 | 1.90 ± 2.35 |
| PI | 0.39 ± 0.22 | 0.69 ± 0.38 | 0.46 ± 0.26 |
| PG | 0.01 ± 0.02 | 0.07 ± 0.08 | 0.04 ± 0.05 |
| LPC | 1.73 ± 1.12 | 2.18 ± 1.59 | 2.70 ± 1.97 |
| LPE | 0.16 ± 0.08 | 0.21 ± 0.14 | 0.20 ± 0.15 |
| LPI | 0.15 ± 0.13 | 0.15 ± 0.19 | 0.16 ± 0.12 |
| LPA | 0.18 ± 0.13 | 0.22 ± 0.20 | 0.18 ± 0.13 |
| AEA | 0.01 ± 0.01 | 0.01 ± 0.02 | 0.03 ± 0.02 |
| SM | 1.45 ± 0.71 | 1.03 ± 0.72 | 0.57 ± 0.80 |
| SPH | 0.08 ± 0.10 | 0.13 ± 0.21 | 0.18 ± 0.27 |
| Cer | 0.49 ± 0.44 | 0.28 ± 0.13 | 0.18 ± 0.15 |
| TG | 27.39 ± 8.84 | 23.58 ± 7.87 | 39.20 ± 13.37 |
| DG | 23.49 ± 11.57 | 28.04 ± 19.43 | 26.52 ± 19.74 |
| MG | 0.04 ± 0.09 | 0.16 ± 0.12 | 0.03 ± 0.06 |
| AcCa | 0.15 ± 0.05 | 0.22 ± 0.25 | 0.32 ± 0.24 |
| Co | 0.04 ± 0.03 | 0.03 ± 0.04 | 0.01 ± 0.02 |

Species-resolved lipid analysis of whole tau-treated versus control organoids revealed modest changes across lipid species (Fig. S2d), reflecting the combined contributions of distinct cellular states, tau_A647_+ and tau_A647_-, within the organoid. To resolve cell state-specific effects, we compared sorted tau-internalizing, tau_A647_+ and tau-exposed but non-internalizing, tau_A647_-, populations. This analysis confirmed a selective reorganization of TG species (Fig. 2g). A large fraction of TG molecules exhibited negative log₂ fold changes in tau_A647_+ cells, indicative of coordinated TG depletion relative to tau_A647_- populations. In contrast, phospholipid and sphingolipid species showed no systematic remodeling. Most PC species, along with subsets of PE, phosphatidylinositol (PI), phosphatidylglycerol (PG), lysophospholipids, and ceramides, displayed only modest positive fold changes. These species-level alterations support a direct association between tau internalization and mobilization of TG-rich lipid stores.

To further characterize this remodeling, we examined TG composition by total acyl carbon number. TG depletion in tau_A647_+ cells was selective for species containing >48 total acyl carbons across the three FA chains (Fig. 2h). Long-chain TGs are more hydrophobic and preferentially stored in LDs ^83^, supporting that these constitute the major fraction of mobilized lipids. Medium- and long-chain FAs (MCFAs and LCFAs) are also preferred substrates for mitochondrial lipid oxidation, raising the possibility that reduced abundance of MCFA- and LCFA-containing TG species in tau_A647_+ cells reflect not only neutral lipid mobilization but also increased utilization through β-oxidative pathways. While acylcarnitine levels (Fig. 2g) did not indicate a global increase in FA transport into mitochondria, localized or transient lipid oxidation cannot be excluded. Analysis of lipid saturation further revealed a selective depletion of polyunsaturated TG species (1-5 double bonds) in tau_A647_+ cells, while tau_A647_- cells showed accumulation of saturated and unsaturated TGs (Fig. 2i). This data suggests that tau-associated lipid remodeling selectively impacts the levels of unsaturated neutral lipid species between tau-internalizing and surrounding, non-internalizing cell populations. Together, integration of the lipidomic and transcriptomic datasets supports a model in which spreading tau drives selective mobilization of neutral lipids and redistribution of long-chain PUFAs between neural cell populations.

### Spreading tau selectively associates with intracellular neutral lipid compartments

The lipid metabolic changes observed in tau-spreading cells prompted us to examine whether tau directly associates with intracellular neutral lipid structures during spreading. Owing to its intrinsically disordered and amphipathic nature, tau can transiently bind lipids ^84–88^. Coupled with the pronounced remodeling of neutral lipid pools, this interaction may influence tau internalization, intracellular localization as well as intercellular lipid communication.

To test this, we performed live-cell imaging of tau_A647_-conditioned iPSC-derived forebrain neurons co-stained with BDP or LipidSpot, an alternative neutral lipid dye. Internalized tau formed discrete intracellular puncta, a subset of which associated with neutral lipid compartments (Fig. 3a,b and Fig. S3b and Table 2). Spot detection and coordinate-based colocalization analysis (Fig. S3a) showed that 21.8 ± 5.5% of tau puncta overlapped with BDP-positive structures at 24 h, increasing to 28.3 ± 9.1% by 48 h (Fig. 3b). Reciprocally, 21.0 ± 6.7% of BDP-positive structures colocalized with tau puncta at 24h, increasing to 26.7 ± 9.3% by 48h (Fig. 3c). Consistent with this, 15.0 ± 4.9% of tau puncta colocalized with the LD coat protein PLIN2 (Fig. S3c and Table 2). Similar tau-lipid associations were detected in mouse primary cortical neurons (Fig. S3d and Table 2), suggesting that lipid localization of spreading tau is conserved across species and neuronal maturation states. To determine the structural basis of this interaction, we compared tau isoforms and domain variants. Lipid association was comparable between the 1N4R and full-length 2N4R isoforms, whereas a C-terminally truncated tau variant (1-245), lacking the microtubule-binding region and extended C-terminus (dMTBR), failed to efficiently internalize or associate with neutral lipids (Fig. S3e, f).

**Figure 3.**
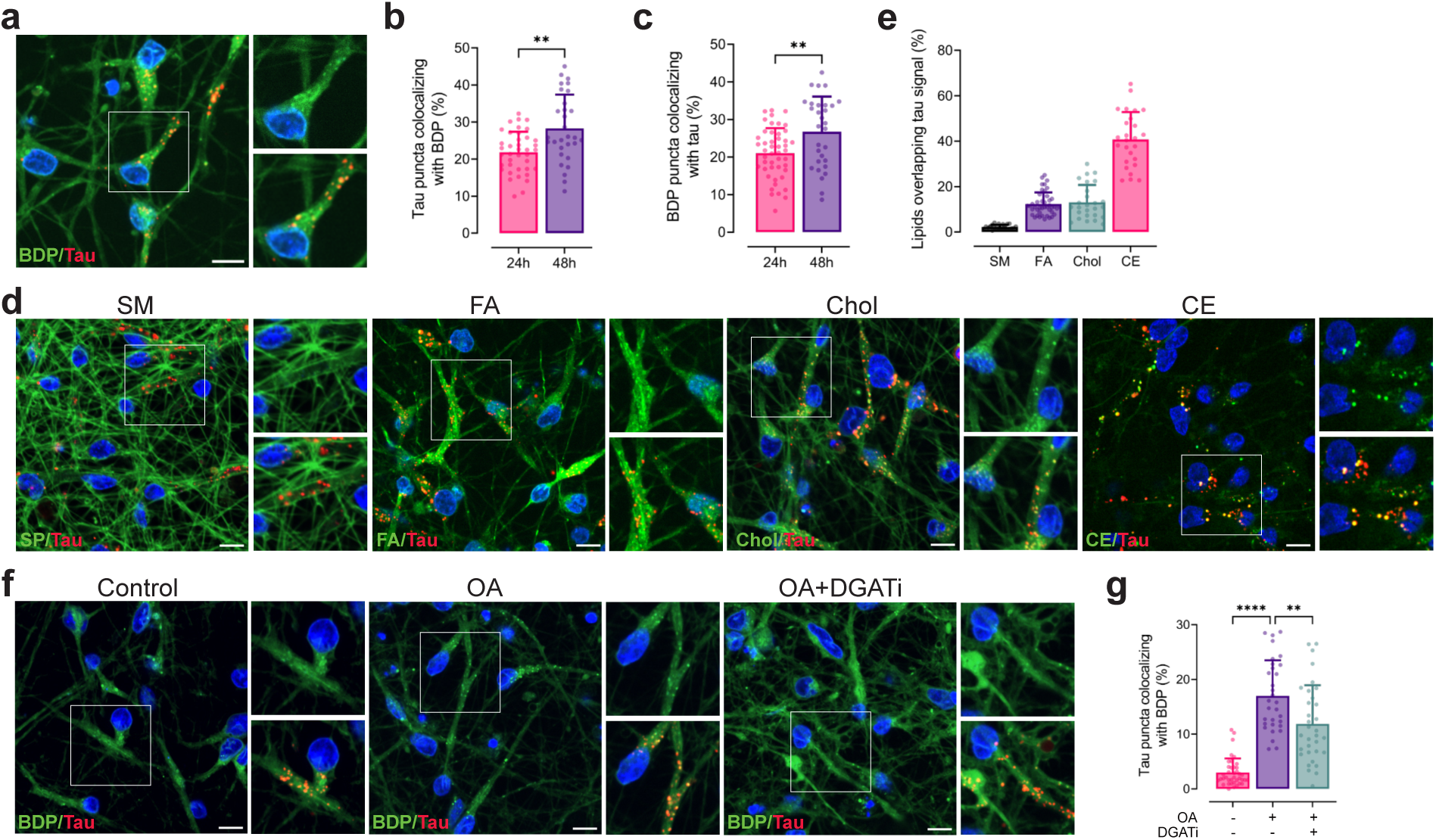
Spreading tau selectively associates with intracellular neutral lipids in neurons. (a) Live-cell imaging of 24 h tau_A647_ (in red)-conditioned iPSC-derived forebrain neurons stained with the neutral lipid dye BDP (green). Insets show tau-BDP colocalization at higher magnification. (b-c) Coordinate- based colocalization. (b) Percentage of tau puncta colocalizing with neutral lipid puncta at 24 h and 48 h after tau exposure (*p* = 0.0013). (c) BDP puncta fraction colocalizing with tau_A647_ at 24 h and 48 h following tau exposure (*p* = 0.0053). In (b-c), *n* = 39 (24h) and *n* = 30 (48h) FOVs, 3 independent experiments. Data were analyzed by Welch’s t-test. (d) Live-cell imaging of neurons treated with fluorescent lipid analogues: sphingomyelin (SM), fatty acid (FA), cholesterol (Chol), and cholesteryl ester (CE) (green), together with tau_A647_ (red). Insets show examples of colocalization or lack thereof. (e) Intensity-based Mander’s coefficient colocalization of tau with the indicated lipid classes at 24 h, displaying the percentage of lipid signal overlapping tau signal: SM 2.4 ± 0.9 %, FA 12.3 ± 5.1 %, Chol 13.1 ± 7.6 % and CE 40.7 ± 12.1 % of lipid overlapping tau signal. *n* = 26 (SM, Chol, CE) and *n* = 46 (FA) FOVs, 3 experimental replicates. (f) Live- cell imaging of neurons conditioned with tau_A647_ (red), OA-treated neurons (+ OA - DGATi), and OA-treated neurons in the presence of DGAT1/2 inhibitor (+ OA + DGATi), all stained with BDP (green). Insets show tau-BDP colocalization. (g) Quantification of coordinate-based colocalization of tau and BDP puncta upon neutral lipid modulation with OA and DGATi. *n* = 39 (- OA - DGATi), *n* = 31 (+ OA - DGATi) and *n* = 33 (+ OA + DGATi) FOVs, 3 independent experiments. Data were analyzed by one-way ANOVA followed by a post-hoc Tukey’s test was used to determine P value. For all images, nuclei are blue. Scale bars, 10 µm.

**Table 2.** Summary % of tau colocalization with intracellular markers.

| Organism | Markers | Method | % |
| --- | --- | --- | --- |
| Human | BDP | Coordinate | 21.8 ± 5.5 |
| Human | LipidSpot | Coordinate | 23.9 ± 11.6 |
| Mouse | BDP | Coordinate | 24.1 ± 11.0 |
| Human | PLIN2 | Coordinate | 15.0 ± 4.9 |

To determine the specificity of tau association with lipids, we performed live-cell imaging of iPSC-derived forebrain neurons using different fluorescent lipid analogs representing major lipid classes. The panel included a neutral lipid, CE; two precursor species capable of esterification to neutral lipids, FA and cholesterol (Chol); and a membrane-associated lipid, sphingomyelin (SM) as a control. Unlike BDP staining, which labels endogenous neutral lipid stores, fluorescent lipid analogs track the intracellular localization and accumulation of defined lipid species. Spreading tau_A647_ preferentially colocalized with intracellular neutral lipids, like CEs, representing 40.7 ± 12.1%, whereas membrane-bound SMs exhibited minimal spatial overlap with tau (2.2 ± 0.9%, Fig. 3d, e). In contrast, FA and Chol analogs displayed predominantly diffuse intracellular distributions, with only a subset of the signal accumulating in discrete puncta. Tau showed partial colocalization with FA- and Chol-positive punctate structures (12.3 ± 5.1% and 13.1 ± 7.6%, respectively; Fig. 3d, e). These puncta likely correspond to newly esterified neutral lipid pools (TGs and CEs) synthetized from the added fluorescent precursor FAs and Chol. This suggests that tau preferentially associates with storage lipid assemblies. Notably, tau-FA colocalization increased over time (Fig. S3g h), consistent with progressive esterification of FA analogs into intracellular neutral lipid-rich compartments.

To test whether elevating intracellular neutral lipids enhances tau interaction with lipids, we pharmacologically modulated TG synthesis. Supplementation with oleic acid (OA) increased the abundance of BDP⁺ LDs by approximately ten-fold (Fig. 3f and S3i) and significantly enhanced tau-lipid colocalization (Ctrl: 3.0 ± 2.5% vs. OA: 17.0 ± 6.5% tau puncta colocalizing with BDP, *p* < 0.0001, Fig. 3f, g)). Conversely, inhibition of TG synthesis using diacylglycerol acyltransferase 1 and 2 (DGAT1 and 2) inhibitors (DGATi) during OA treatment markedly reduced LD abundance (Fig. 3f and S3i) and attenuated tau-lipid colocalization (OA: 17.0 ± 6.5% vs. OA+DGATi: 11.9 ± 7.0 %% tau puncta colocalizing with BDP, *p* = 0.0011, Fig. 3f, g). Together, these results demonstrate that the association of spreading tau with intracellular neutral lipid species is metabolically regulated, requiring active lipid esterification, and is enhanced by elevated neuronal lipid content.

### Spreading tau forms dynamic, neuron-specific lipid assemblies with distinct structural states

Neutral lipid metabolism in the brain is shaped by metabolic coupling between neurons and astrocytes. While astrocytes exhibit a greater lipid storage capacity, neurons have high metabolic demands but limited capacity to store lipids in LDs. Nevertheless, recent work demonstrated that neurons not only rely in glucose catabolism, but also actively engage in TG turnover to support bioenergetic demands ^89,90^. We next asked whether tau-lipid association depends on cell identity and intrinsic lipid metabolic states. Internalized tau_A647_ colocalized with neutral lipids in neurons (shown in Fig. 3a) but showed minimal overlap with LDs in astrocytes, where tau puncta frequently localized adjacent to, rather than within, neutral lipid structures (N (Neurons): 22.1 ± 5.5 % vs. A (Astrocytes): 4.2 ± 2.6 % of tau puncta colocalizing with BDP, *p* < 0.0001, Fig. 4a,b). In SH-SY5Y cells, a commonly used human neuroblastoma-derived cell line, tau internalized but failed to associate with neutral LDs (Fig. S4a). This suggests that tau-lipid association is not a general consequence of uptake and may depend on neuron-specific intracellular trafficking or post-endocytic processing states that might direct tau to adopt a distinct structural state.

**Figure 4.**
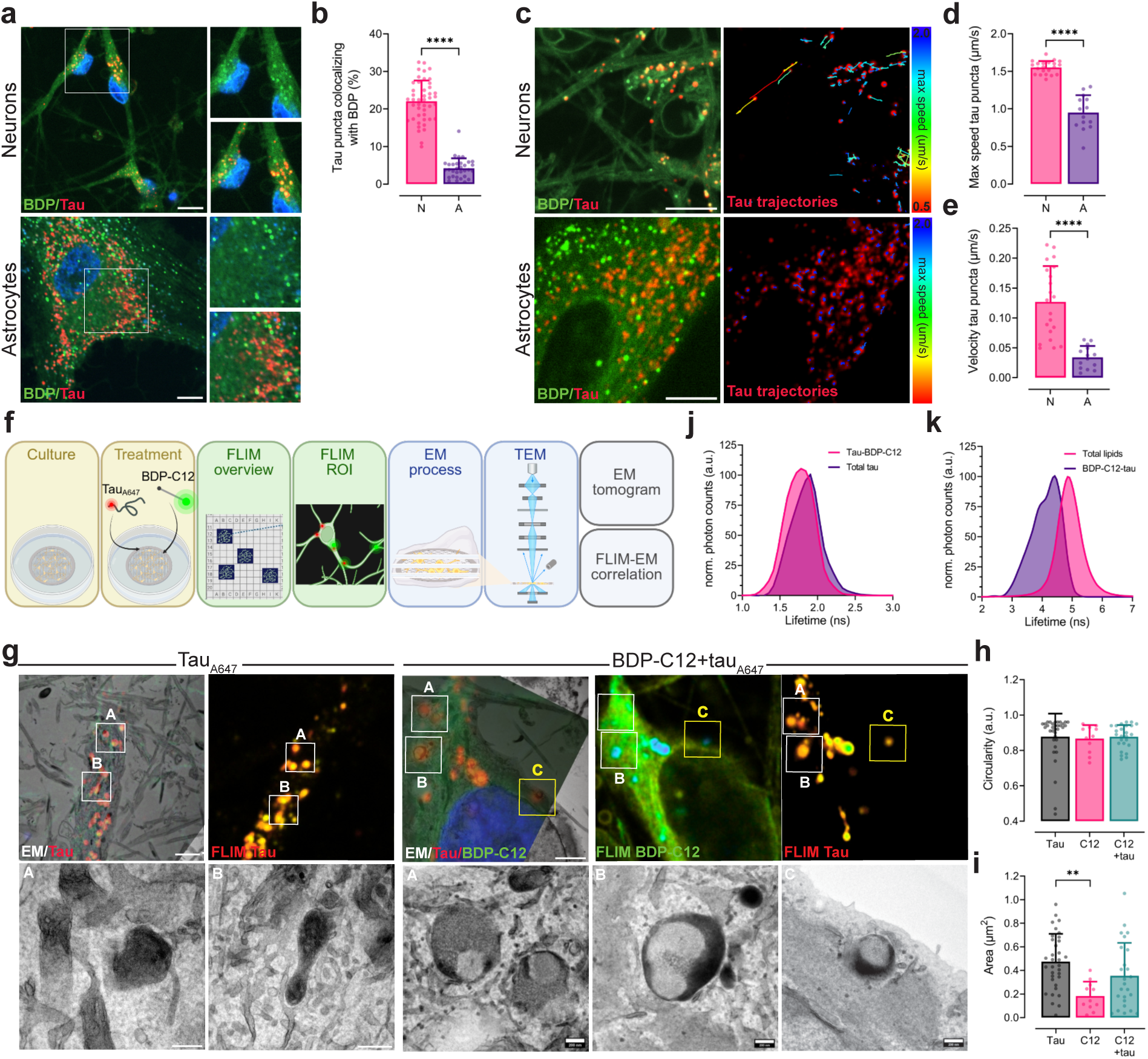
Spreading tau forms neuron-specific, dynamic lipid-associated assemblies with distinct structural states. (a) Live-cell imaging of iPSC-derived neurons (top) and astrocytes (bottom) conditioned with tau_A647_ (red) and BDP (green). Insets show examples of tau puncta colocalizing with BDP. Nuclei are in blue. Scale bars, 10 µm. (b) Coordinate-based colocalization analysis displaying the percentage of tau_A647_ puncta colocalizing with BDP-positive structures in neurons and astrocytes. *n* = 45 FOVs (neurons), and *n* = 32 FOVs (astrocytes),3 independent experiments. Statistical significance was determined using Welch’s test. (c) Single-particle tracking of tau_A647_ puncta in neurons (top) and astrocytes (bottom). Left: Representative frame of time-lapse measurement of tau_A647_(red)- conditioned cells co-stained with BDP (green). BDP signal was used to identify cell morphology. Right: color-coded trajectories of individual tau puncta. The color scale ranges from red (minimum, 0.5 µm/s) to blue (maximum, 2 µm/s) track maximum speed. (d-e) Quantification of tau puncta dynamics. (d) Maximum speed and (e) mean velocity of tau puncta in neurons and astrocytes. *n* = 21 neuronal and *n* = 14 astrocytic time-lapse recordings,4 and 3 independent experiments, respectively. P values were obtained by analyzing the data using Welch’s test. (f) Correlative FLIM–EM workflow in neurons following 48 h exposure to BDP-C12, tau_A647_, or combined BDP-C12 and tau_A647_ treatment. (g) Left (Tau): Representative correlated FLIM-EM images of iPSC-derived neurons conditioned with tau_A647_ (red): Top left: overview FLIM-EM correlation. Boxed areas (A and B) highlight the ROIs shown at the bottom. Scale bars, 2 µm. Top right: corresponding FLIM image of tau_A647_. Color-scale bar indicates the range of lifetimes (ns). Bottom: electron micrographs of highlighted ROIs (A and B). Scale bars, 200 nm. Right (BDP-C12+tau): Representative correlated FLIM-EM images of neurons conditioned with tau_A647_ (red) and BDP-12 (green); Top left: overview of FLIM-EM correlation. Boxed areas (A, B and C) highlight the electron micrographs shown at the bottom. Scale bars, 2 µm. Top center: corresponding FLIM image of BDP-C12: Top right: corresponding FLIM image of tau_A647_. Color-scale bars indicates the range of lifetimes (ns). Bottom: electron micrographs of highlighted ROIs (A, B and C). Scale bars, 200 nm. The tomogram of ROI C is shown in Fig. SI4i. BDP-C12 1 experimental replicate, tau_A647_ and combined BDP-C12 and tau_A647_ 3 experimental replicates. (h) Quantification of the circularity of puncta structures identified by FLIM-EM correlation: Tau 0.88 ± 0.13, BDP-C12 0.87 ± 0.07 and BDP-C12+tau 0.88 ± 0.07 a.u., *n* = 34 (Tau), *n* = 11 (BDP-C12) and *n* = 24 (BDP-C12+tau) individual puncta, 2 experimental replicates. P value was calculated by Kruskal-Wallis test. (i) Size puncta quantification of the structures identified by FLIM-EM correlation: Tau 0.47 ± 0.23, BDP-C12 0.18 ± 0.12 and BDP-C12+tau 0.35 ± 0.27 a.u., *p* = 0.0024. *n* = 34 (Tau), *n* = 11 (BDP-C12) and *n* = 24 (BDP-C12+tau) individual puncta, 2 experimental replicates. P value was calculated by one-way ANOVA followed by a Tukey’s multiple comparison test. (j) Fluorescence lifetime distribution of tau_A647_ assemblies in neurons under lipid-enriched and non-enriched conditions. (k) Fluorescence lifetime distribution of BDP-C12-positive structures colocalizing with tau puncta compared with the total BDP-C12 population in neurons.

Live-cell imaging and single-particle tracking revealed that neuronal tau-lipid assemblies exhibited rapid, directional motion (maximum speed = 1.55 ± 0.08 µm s⁻¹; velocity = 0.12 ± 0.06 µm s⁻¹), whereas tau puncta in astrocytes remained comparatively static (maximum speed = 0.94 ± 0.23 µm s⁻¹, *p* < 0.0001; velocity = 0.03 ± 0.02 µm s⁻¹, *p* < 0.0001, Fig. 4c-e and movie 3 and 4). The dynamic behavior of tau assemblies was further confirmed by monitoring fusion and fission using sequential conditioning with tau_A594_ and tau_A647_. Neuronal tau assemblies underwent frequent fission and fusion events, with tau_A594_ and tau_A647_ signals merging over time, whereas in astrocytes, the two populations remained largely distinct (Fig. S4b, c and movie 5 and 6). Treatment with 1,6-hexanediol (10% w/v), an aliphatic alcohol that weakens hydrophobic interactions and dissolves liquid-liquid phase separated droplets^91^, partially reduced the size of tau-lipid assemblies in neurons (Fig. S4d, e). These observations suggest that neuronal tau-lipid assemblies are maintained by weak, hydrophobic interactions and display fluid, coalescent properties. The rapid and directional motion of neuronal tau-lipid assemblies suggested active transport along the cytoskeleton. Given the established association of endogenous tau with microtubules^79^, we next examined whether spreading tau similarly engages the microtubule network. Live imaging of neurons stained for tubulin revealed that tau_A594_ puncta dynamically translocated along microtubule tracks, with 82.5 ± 11.2% of trajectories overlapping with tubulin signal (Fig. S4f and movie 7). In contrast, overexpressed SNAP-tagged tau localized on microtubules, as expected, but showed minimal colocalization with neutral lipids (Fig. S4g). This divergence indicates that exogenously expressed tau and internalized spreading tau assemblies represent distinct molecular states with different subcellular localization and intracellular interactions partners.

To resolve the ultrastructural organization of these tau assemblies in neurons, we established a correlative fluorescence lifetime imaging microscopy and electron microscopy (FLIM-EM) pipeline. Following 48h of tau_A647_ treatment, FLIM imaging on neurons was performed prior to EM processing (Fig. 4f). This workflow preserved the ultrastructural integrity of low-abundance tau puncta and mitigated the effects of fluorescence loss on detection. Using this approach, we observed membrane-associated tau structures with morphologies similar to those detected in organoids (Fig. 1e). Correlative EM revealed that tau_A647_ inclusions were predominantly spherical and localized within membrane-bound compartments, with an average cross-sectional area of 0.47 µm² (± 0.27 SD; *n* = 31 correlated tau structures; Fig. 4g-i). Neurons enriched with the fluorescent FA analog (BDP-C12) contained membrane-bound lipid compartments with similar circularity but significantly smaller size than tau-containing structures (Fig. Fig. 4h-i and S4h). Following tau_A647_ treatment, we detected tau-lipid rich assemblies in which amorphous tau oligomers frequently extended from the limiting membrane of these compartments toward their interior. The central lumen remained electron-lucent and occasionally contained smaller vesicles or electron-dense lipid material, a morphology consistent with multivesicular bodies or endosomal compartments. Notably, tau and lipid-rich domains occupied spatially distinct regions within the same compartment, giving rise to a biphasic organization that was conserved between neurons and organoids (Fig. 4g and Fig. 1e). Quantitative analysis further showed that tau-associated lipid compartments were larger than BDP-C12-positive compartments lacking tau (Fig. 4i). Electron tomography further revealed contacts with the endoplasmic reticulum, plasma membrane, and many of these compartments were positioned adjacent to microtubules (Fig. S4i and movie 8). Moreover, the FLIM-EM correlative approach allowed us to investigate changes in the microenvironment of tau-lipid rich assemblies. Tau_A647_ fluorescence lifetime distribution differed between lipid-enriched and non-enriched conditions (LT = 1.78 ns vs. 1.90 ns; Fig. 4g,j), indicating that lipid loading alters the local molecular environment surrounding tau assemblies. Tau-associated lipids exhibit fluorescence lifetime distributions distinct from the bulk cellular BDP-C12 pool, indicating that spreading tau resides within a biochemically distinct neutral lipid microenvironment (total lipids: 4.84 ns vs. lipids associated with tau: 4.42 ns; Fig. 4g,k). Together, these findings indicate that spreading tau forms dynamic, lipid-rich intracellular assemblies with distinct molecular conformation that undergo microtubule-associated trafficking in neurons.

### Spreading of tau in neurons promotes lipid clearance by facilitating lipid transfer to astrocytes

Neurons rely on multiple mechanisms to maintain lipid homeostasis and prevent lipotoxicity, including lysosomal degradation and intercellular lipid transfer to astrocytes^50^. Particularly during periods of high neuronal activity, neurons offload excess FAs, including peroxidated species, to astrocytes, which store them in LDs, detoxify or metabolize them, thereby protecting neurons from lipotoxic damage ^92^. Disruptions in this neuronal to astrocyte lipid flux have been implicated in tauopathies ^59,93^ Given our findings that spreading tau associates with, and remodels the pools of intracellular neutral lipids, we hypothesized that spreading tau may influence lipid clearance in neurons.

To test our hypothesis, we used an established neuron-astrocyte co-culture lipid transfer assay ^50^. Neurons were incubated with BDP-C12, with or without spreading tau_A647._ Next, neurons were co-cultured with astrocytes in non-contact conditions (Fig. 5a). Following 4h of co-culture, tau-conditioned neurons were able to clear excess FA more rapidly than control neurons, exhibiting an approximately 50% reduction in BDP- C12 intensity (Ctrl; 0.60 ± 0.21 vs. Tau; 0.33 ± 0.22; *p* < 0.0001, Fig. 5b, c). This enhanced lipid clearance was accompanied by increased accumulation of neuronal BDP-C12 signal in astrocytes when co-cultured with tau-conditioned neurons (Fig. S5a, b). After 24 h of co-culture, elevated BDP-C12 signal persisted in astrocytes in tau-conditioned co-cultures (Ctrl; 0.30 ± 0.20 vs. Tau; 0.59 ± 0.24; *p* < 0.0001, Fig. 5d, e), and neuronal tau_A647_ was detectable in astrocytes (Fig. 5d, f), consistent with continued transfer of both lipid cargo and spreading tau between cell types.

**Figure 5.**
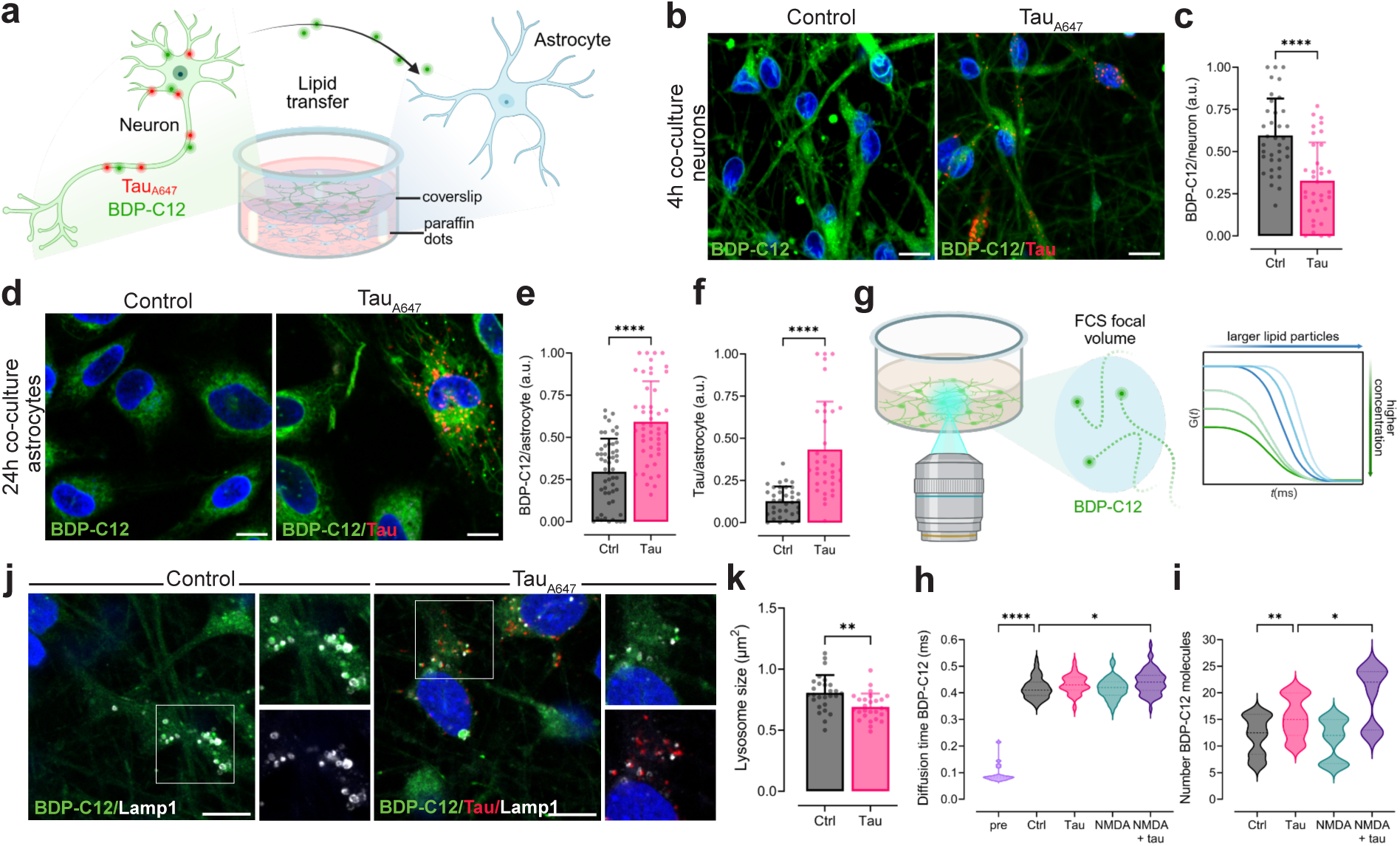
Spreading tau enhances neuronal lipid efflux and transfer to astrocytes while relieving lysosomal lipid stress. (a) Schematic of the co-culture setup used to monitor neuron-to-astrocyte lipid transfer (b) Representative images of BDP-C12 signal (green) in neurons after 4h of co-culture comparing tau_A647_ (red)-conditioned with control neurons. (c) Quantification of BDP-C12 intensity per neuron in control and tau exposed neurons, *n* = 36 (Ctrl) and *n* = 35 (Tau) FOVs, 3 independent experiments. (d) Representative images of astrocytes co- cultured for 24 h with control or tau_A647_-conditioned neurons, showing BDP-C12 (green) and tau_A647_ (red). (e) Quantification of BDP-C12 signal (green) in astrocytes after 24h of co-culture, *n* = 53 (Ctrl) and *n* = 48 (Tau) FOVs, 4 experimental replicates. (f) Quantification of tau_A647_ intensity per astrocyte after 24 h of co- culture with neurons: Ctrl: 0.13 ± 0.09 vs. Tau: 0.43 ± 0.28 (a.u.), *n* = 36 (Ctrl) and *n* = 35 (Tau) FOVs, 3 experimental replicates. For (c, e and f) p value was calculated using a two-tailed unpaired t-test. (g) Schematic of FCS to detect secreted lipid particles into the media (h) Diffusion time of BDP-C12 particles measure by FCS compared across conditions: before incubation with neurons (pre: 0.09 ± 0.03 ms), control (Ctrl: 0.42 ± 0.04 ms), tau_A647_ (Tau: 0.43 ± 0.03 ms), NMDA (NMDA: 0.41 ± 0.04) and NMDA + tau_A647_ (NMDA=tau: 0.44 ± 0.04 ms); pre vs. Ctrl *p* < 0.0001 and Ctrl vs. NMDA+tau *p* = 0.0488; *n* = 60 (ctrl and NMDA) and *n* = 59 (tau and NMDA+tau), 3 independent experiments. (i) Number of BDP-C12-positive particles detected by FCS in media from control (Ctrl: 12 ± 3 molecules), tau_A647_-treated (Tau: 15 ± 4 molecules), stimulated with NMDA (NMDA: 11 ± 4 molecules) or NMDA-stimulated exposed to tau_A647_ (NMDA+tau: 20 ± 5 molecules) in mono-cultured neurons; Ctrl vs. Tau *p* = 0.042 and Tau vs. NMDA+tau *p* = 0.0117; *n* = 60 (ctrl and NMDA) and *n* = 59 (tau and NMDA+tau) from 3 independent experiments. For (h and i), P values were calculated using a Kruskal-Wallis test follow by a post-hoc Dunn’s multiple comparisons test. (j) Lamp1 staining (lysosomes, grey scale) in control and tau_A647_ (red)-treated neurons both supplemented with BDP-C12 (green); insets show enlarged views of Lamp1-positive structures. (k) Quantification of lysosome size: Ctrl: 0.81 ± 0.15 µm^2^ vs. Tau: 0.69 ± 0.11 µm^2^; p = 0.0019; *n* = 27 FOVs for both conditions, 3 independent experiments. P value was determined by two-tailed unpaired t-test. Nuclei are in blue. Scale bars, 10 µm.

Next, we asked whether tau-mediated FA clearance is a neuron-intrinsic mechanism or depends on astrocytic ApoE. We used fluorescence correlation spectroscopy (FCS), a single-molecule sensitive approach to detect fluorescent lipid particles released into the medium of mono-cultured neurons, to directly quantify neuronal lipid efflux (Fig. 5g). Diffusion times of extracellular fluorescent species were substantially longer than those of free BDP-C12 in culture medium (Fig. 5h), indicating that the detected signal represents lipids that had been internalized, metabolically processed, and subsequently secreted as larger lipid-containing particles. We found that neurons exposed to spreading tau consistently secreted more lipid particles into the extracellular space compared to control neurons (Fig. 5i). Combined tau and N-methyl-D-aspartate (NMDA) treatment, which induces neuronal hyperactivity and secondary oxidative damage^94,95^, further amplified lipid secretion, with neurons releasing a higher number and slightly larger particles compared to tau alone (Fig. 5h, i), suggesting that increased neuronal activity or the resulting oxidative stress amplifies tau-dependent lipid export. Together, these data indicate that spreading tau robustly promotes neuronal lipid efflux in a cell-autonomous manner, independent of astrocyte contact. In contrast to astrocytic ApoE, which functions as a constitutive lipid transporter, neuronal ApoE expression is induced during stress^72^ potentially helping neurons reduce lipotoxicity We next tested whether tau-induced lipid clearance requires neuronal ApoE. In neurons supplemented with BDP-C12, immunostainings revealed a slight reduction in ApoE levels in tau-conditioned neurons (Fig. S5c, d), consistent with transcriptomic data showing APOE downregulation in tau-conditioned organoids (Fig. 2b). Despite this decrease, neuronal lipid clearance was enhanced, demonstrating that spreading tau may engage an ApoE-independent efflux mechanism.

Lysosomal degradation of accumulating lipids represents an alternative neuronal mechanism for preventing lipotoxicity. Lysosomes accumulating oxidized lipids become swollen. Thus, lysosomal enlargement has been linked to lipid metabolic stress^96^.. We next analyzed lysosomal morphology via Lamp1 immunostaining in BDP-C12 treated neurons comparing tau-conditioned neurons and controls. Quantitative analysis revealed enlarged Lamp1-positive structures in neurons accumulating BDP-C12, whereas tau-conditioned neurons displayed smaller lysosomes (Fig. 5j, k). This is consistent with the reduced BDP-C12 burden observed in tau-conditioned neurons (Fig. 5b, c), together supporting a role for tau-driven lipid clearance in reducing lysosomal lipid burden and maintaining neuronal lipid homeostasis.

### Tau spread enhances neuronal resilience to oxidative stress through lipid detoxification mechanisms

Lipidomic analyses indicated an accumulation of unsaturated lipid species in tau_A647_- cells (Fig. 2i), a class of lipids particularly vulnerable to peroxidation. Consistent with a link to oxidative stress, FCS measurements demonstrated that tau-dependent lipid secretion was amplified under NMDA treatment, which induces oxidative stress (Fig. 5i). We next determined whether spreading tau preferentially associates with oxidized lipids using the ratiometric lipid peroxidation probe BODIPY 581/591 C11 (reduced lipids (C11red), oxidized lipids (C11ox)). Under basal conditions, tau puncta showed a strong preference for oxidized lipid assemblies over non-oxidized lipid pools in both human iPSC-derived forebrain neurons (Fig. 6a, b) and primary mouse cortical neurons (Fig. S6a, b). More than 25% of internalized tau_A647_ localized to sites of lipid peroxidation (Fig. 6c). To modulate peroxidized lipids levels, neurons were treated with NMDA, which produced the expected increase in C11ox intensity (Fig. S6c, d). We next tested weather elevated peroxidized lipid levels following NMDA stimulation influence tau recruitment. Indeed, we detected enhanced tau recruitment to C11ox-positive lipid compartments, without altering its localization to reduced lipid pools (Fig. 6c), identifying intracellular neutral lipids rich in oxidized species as the major intracellular destination of spreading tau. To determine whether astrocytes participate in the clearance of tau-associated oxidized lipids, NMDA- treated neurons were co-cultured with astrocytes. Under conditions of elevated lipid peroxidation, tau conditioning significantly reduced neuronal C11ox levels in co-culture conditions (Fig. 6d, e). While neurons retained higher oxidized lipid burden in monoculture (NMDA+tau: 0.42 ± 0.25), co-culture markedly attenuated neuronal C11ox signal (0.20 ± 0.19; *p* < 0.0001; Fig. 6e). In parallel, astrocytic C11 ox/red ratio increased in tau-conditioned co-cultures compared with NMDA-treated controls (Fig. 6f, g), consistent with transfer of lipid peroxides from neurons to astrocytes. To determine whether this mechanism extends beyond excitotoxic stress, ferroptosis was induced with erastin, which disrupts glutathione metabolism and drives direct iron-dependent lipid peroxidation^97^. Erastin treatment, as expected, elevated neuronal C11ox (Fig. S6e, f). Under ferroptotic conditions induced by erastin treatment tau enhanced peroxidized lipid clearance in neurons co-cultured with astrocytes (Fig. S6g, h) while increased the astrocytic C11ox/C11red ratio (Fig. S6i, j), indicating that tau-mediated lipid detoxification operates across distinct oxidative stress paradigms. These results identify tau spreading as a mediator of neuron-to-astrocyte transfer of peroxidized lipids that is required for effective neuronal detoxification across multiple oxidative stress conditions.

**Figure 6.**
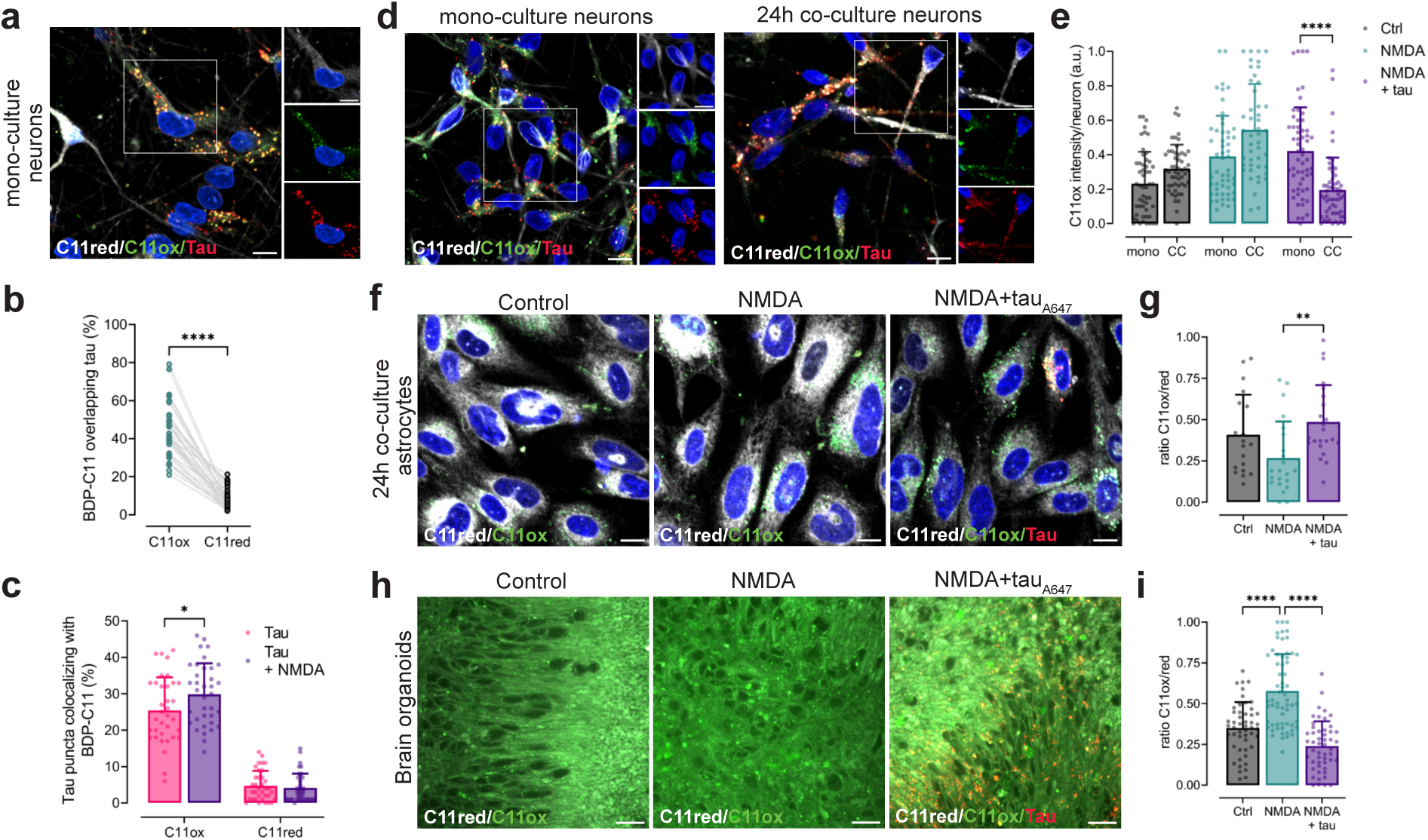
Spreading tau targets oxidized lipids and promotes astrocyte-mediated clearance of lipid peroxides. (a) Live-cell imaging of human iPSC-derived forebrain neurons labeled with the radiometric lipid peroxidation sensor BDP-C11: C11red, reduced lipids (grey); C11ox, oxidized lipids (green) and tau_A647_ (red). Insets show tau puncta colocalizing with C11ox-positive lipid assemblies. (b) Intensity-based colocalization analysis displaying percentage of tau signal overlapping with C11ox (44 ± 13 %) versus C11red (8 ± 5 %; p < 0.0001)) under basal conditions assessed by Mander’s coefficient, *n* = 47 FOVs, 3 independent experiments. Wilcoxon test was used to assess significance. (c) Coordinate-based colocalization assessing the percentage of tau_A647_ puncta colocalizing with C11ox or C11red puncta in Tau neurons (C11ox: 25 ± 9% and C11red: 5 ± 4 %) and upon NMDA stimulation (C11ox: 30 ± 8 % and C11red: 4 ± 4 %); *n* = 35 (Tau) and *n* = 36 (NMDA+tau) FOVs, 3 independent experiments. Data was analyzed by two-way ANOVA using excitotoxicity (p = 0.0294) and oxidation state (*p* < 0.0001) as independent variables, post-hoc Tukey’s C11ox in tau vs C11ox in tau +NMDA *\*p* = 0.035). (d) Representative images comparing BDP-C11(C11red (grey); C11ox (green)) signal between neurons exposed to tau_A647_ (red) in monoculture (mono, left) compared to neurons in co-culture (CC, right, both under NMDA stress (NMDA+tau). Insets highlight C11ox- tau puncta. (e) Quantification of neuronal C11ox intensity across conditions in (d): Ctrl/mono: 0.23 ± 0.18 a.u. *n* = 52, NMDA/mono: 0.39 ± 0.24 a.u. *n* = 53, NMDA+tau/mono: 0.42 ± 0.25 a.u. *n* = 57, Ctrl/CC 0.32 ± 0.14 a.u. *n* = 52, NMDA/CC 0.54 ± 0.27 a.u. *n* = 46 and NMDA+tau/CC 0.19 ± 0.19 a.u. *n* = 58. Ns represent FOVs from 3 independent mono-culture (mono) and co-culture (CC) experiments. Data was analyzed by two-way ANOVA using culture mode (p = 0.8133) and tau/NMDA condition (*p* < 0.0001) as independent variables, post-hoc Tukey’s: NMDA+tau/mono vs.NMDA+tau/CC *p* < 0.0001. (f) Representative images of astrocytes co-cultured for 24 h with control neurons, NMDA-treated neurons, or NMDA-treated neurons exposed to spreading tau (red), neuronal BDP-C11 signal present in the astrocytes (C11red (grey); C11ox (green)). (g) Quantification of astrocytic C11ox/C11red ratio comparing experimental conditions in (f): Ctrl: 0.41 ± 0.24 a.u. *n* = 19, NMDA: 0.27 ± 0.22 a.u. *n* = 22 and NMDA+tau: 0.48 ± 0.22 a.u. *n* = 23. Ns represent FOVs from 2 experimental replicates. Ordinary one-way ANOVA (p = 0.077) followed by a post-hoc Tukey’s test was used to determine P value: NMDA vs. NMDA +tau *p* = 0.059. Scale bars in (a,d,and f), 10 µm, nuclei are in blue. (h) Live-cell imaging of brain organoid labeled with BDP-C11 (C11red (grey); C11ox (green)) under control, NMDA, and NMDA+tau (red) exposure conditions. Scale bars, 20 μm. (i) Quantification of organoid C11ox/C11red ratio: Ctrl: 0.35 ± 0.16 a.u. *n* = 50, NMDA: 0.58 ± 0.22 a.u. *n* = 61 and NMDA+tau: 0.24 ± 0.15 *n* = 51. Ns represent FOVs from 5 individual brain organoids per condition and 3 independent experiments. Statistical significance was determined using an ordinary one-way ANOVA (*p* < 0.0001) followed by post-hoc Tukey’s test; Ctrl vs NMDA and NMDA vs NMDA +tau, for both p < 0.0001.

To assess tissue-level effects of tau spreading on lipid peroxidation, we quantified the BDP-C11 ox/red ratio in brain organoids subjected to NMDA or erastin treatment. Both treatments markedly increased this ratio, confirming elevated lipid peroxidation (Fig. 6h, i and S6k, l). Remarkably, tau conditioning under these lipid peroxidation burdens significantly reduced the BDP-C11 ox/red ratio in both models (NMDA; 0.58 ± 0.23 vs, tNMDA+tau; 0.24 ± 0.15; *p* < 0.0001) and (Erastin: 0.57 ± 0.14 vs Erastin+tau: 0.46 ± 0.20, *p* <0.0001). Together, these findings identify a protective role for spreading tau in mitigating oxidative lipid stress. By promoting the secretion and astrocyte-mediated clearance of oxidized FAs, spreading tau enhances neuronal resilience to excitotoxic and ferroptotic challenges.

## Discussion

This work demonstrates that spreading tau engages a previously unrecognized metabolic program that remodels neutral lipid pools and redistributes lipid species across neural cell types. These findings uncover a coupling between tau transmission and intercellular lipid flux, revealing a metabolic dimension of tau biology that precedes classical pathological hallmarks. Using brain organoids, a multicellular neural tissue model that recapitulates higher complexity than 2D cultures, we captured early stages of tau spreading and defined its impact on lipid metabolism. Exogenously supplied monomeric tau is taken up and spreads in a time- and HSPG-dependent manner while remaining non-fibrillar. In contrast to previous transcriptomic^69,98,99^ and lipidomic^58,66,100^ studies of tauopathy, which primarily capture late-stage tissue responses, our approach resolves the distinct cellular states generated by tau spreading before pathological hallmarks arise. Tau-internalizing and tau-exposed but tau-negative cells within the same tissue adopt different transcriptional and lipid states, indicating that tau internalization and spread defines a discrete metabolic program that partitions the tissue rather than uniformly perturbing the neural environment. Tau- internalizing cells exhibit activation of lipid trafficking and mobilization pathways and depletion of neutral lipids, and reduced LD abundance. Neighboring tau-negative cells engage inflammatory and stress-associated signaling and accumulate TG-rich lipids. This divergence indicates that tau spreading segregates neural cellular environment into asymmetric metabolic states, establishing coordinated redistribution of lipid resources across cell populations. Such compartmentalization may help to explain why tau spreading occurs in both homeostatic states and in early stages of disease without compromising tissue integrity ^7,28,36–38,101^. These findings also raise the possibility that metabolic state may be an active determinant of tau propagation. Future studies should therefore investigate whether distinct metabolic states define permissive or resistant cellular niches that predict the directionality and efficiency of tau transmission across neural networks.

A defining feature of tau-driven lipid remodeling is the neuron-specific interaction of spreading tau with neutral lipid-rich compartments. In neurons, but not astrocytes, tau accumulates in dynamic, coalescent assemblies that associate selectively with esterified neutral lipid species, undergo directed intracellular transport, and localize to heterogeneous membrane-bound compartments. This behavior is distinct from intracellularly expressed tau, which primarily decorates microtubules and lacks comparable lipid association, supporting the idea that spreading tau represents a distinct molecular state with unique intracellular interactions and trafficking properties. The determinants of this neuronal specificity, whether related to LD biology, membrane composition, or intracellular trafficking pathways, remain to be defined. Dissecting this interplay will be important for understanding how extracellular tau acquires its unique intracellular behavior and whether alterations in lipid composition drive the transition from physiological tau assemblies to pathological aggregation.

Functionally, tau spreading promotes FA efflux and the mobilization of peroxidized lipid species from neurons to astrocytes. This tau-driven metabolic handoff is accompanied by reduced lysosomal lipid burden and attenuation of neuronal lipid peroxidation, indicating that tau spreading contributes to lipid detoxification and maintenance of cellular lipid homeostasis. Notably, this occurs despite reduced neuronal APOE expression, suggesting engagement of lipid export pathways distinct from canonical ApoE-mediated stress responses. Upregulation of SORT1, a multi-ligand sorting receptor involved in lipoprotein uptake and intracellular lipid trafficking^102,103^, further implicates regulated lipid trafficking networks and identifies a potential pathway for tau-associated lipid transport. These findings integrate with emerging models of neuron-astrocyte metabolic coupling, in which neurons rely on astrocytes to buffer excess or damaged lipids^92^. Neurons can export FAs under conditions of metabolic or oxidative stress and utilize lipid flux as part of their adaptive response. Tau-associated lipid assemblies may represent a mechanism to mobilize, package, and transfer neutral lipids during stress, linking intracellular lipid handling to intercellular communication pathways. This raises the possibility that tau spreading contributes to the coordination of lipid metabolism across brain tissue.

Our data suggest that tau spreading participates in a physiological, stress-responsive program that facilitates removal of peroxidized lipids from neurons. However, the functional outcome of this process is likely to depend on its magnitude, timing, and coupling to astrocytic buffering capacity. Under acute or moderate stress, spreading tau may support lipid detoxification by mobilizing neutral lipid-rich assemblies and promoting transfer of damaged lipids to astrocytes. Under sustained stress, aging, or disease-associated metabolic imbalance, this pathway may become saturated or uncoupled, leading to extracellular tau accumulation, persistence of tau-containing lipid assemblies, and/or impaired lipid clearance. Recent work demonstrates that tau-lipid interactions are strongly influenced by lipid composition including cholesterol content ^104^. Together with our findings, this suggests that disease-associated lipid remodeling may alter tau recruitment to lipid compartments and disrupt its physiological role in lipid clearance. Post-translational modifications, isoform composition, and conformational changes may further shift tau from a dynamic lipid-associated species toward aggregation-prone assemblies. As such, neutral lipid-rich compartments may represent sites where protective lipid handling and pathological tau accumulation intersect.

We thus propose that tau spreading serves pre-pathogenic, homeostatic functions, that require tight regulation of neuron-astrocyte metabolic communication. Failure, saturation, or biochemical remodeling of these adaptive pathways may contribute to lipid dyshomeostasis, tau accumulation, and neurodegeneration. Indeed, neuronal tau pathology induces accumulation of LDs and unsaturated lipids in microglia^59^, further supporting that tau accumulation intersects with intercellular metabolic communication and brain lipid metabolism homeostasis. Consistent with this model, resilient AD brains exhibit reduced TG levels and fewer LDs^105^ suggesting that efficient neutral lipid turnover may be a hallmark of resilience. Understanding how tau coordinates lipid homeostasis may therefore reveal new therapeutic strategies aimed at preserving endogenous resilience mechanisms, rather than simply preventing tau aggregation, in AD and related tauopathies.

Importantly, tau is not unique in exhibiting extracellular spreading. Other IDPs, including α-synuclein and TDP-43, are also released into extracellular fluids and propagate between cells ^106–108^ Our findings raise the possibility that extracellular IDPs function more broadly as regulators of intercellular metabolic communication instead of acting solely as agents of pathology. The intrinsic structural plasticity of IDPs may be particularly well suited for this extracellular role, enabling dynamic interactions with diverse proteins, membranes, and lipids that can rapidly adapt to changing tissue environments. In this view, neurodegeneration may arise from failure to maintain the balance of extracellular protein-mediated metabolic communication. Elucidating how these processes are regulated, and how they become disrupted, may define new strategies to restore metabolic homeostasis and modify disease trajectories.

## Abbreviations: (non-standard and used 3 or more times)

tau_A647_: 
Tf_A594_: 
tau_A488_: 
BDP: 
BDP-C11: 
BDP-C12: 
C11ox: 
C11red: 
DGATi: 
FLIM-EM: 
LipidSpot: 
ThioS: 
dMTBR: 

---

**Movie 1.** Time-lapse recording of the calcium imaging (CalBryte 520, grey-scale) measurement in organoids showed in Fig. SI1a

**Movie 2.** Time-lapse recording shown in Fig. 1d used to calculate tau_A647_ (red) trajectories by single-particle tracking of tau_A647_ puncta. CalBryte 520 (grey-scale) was used to identify cell morphology.

**Figure S1.**
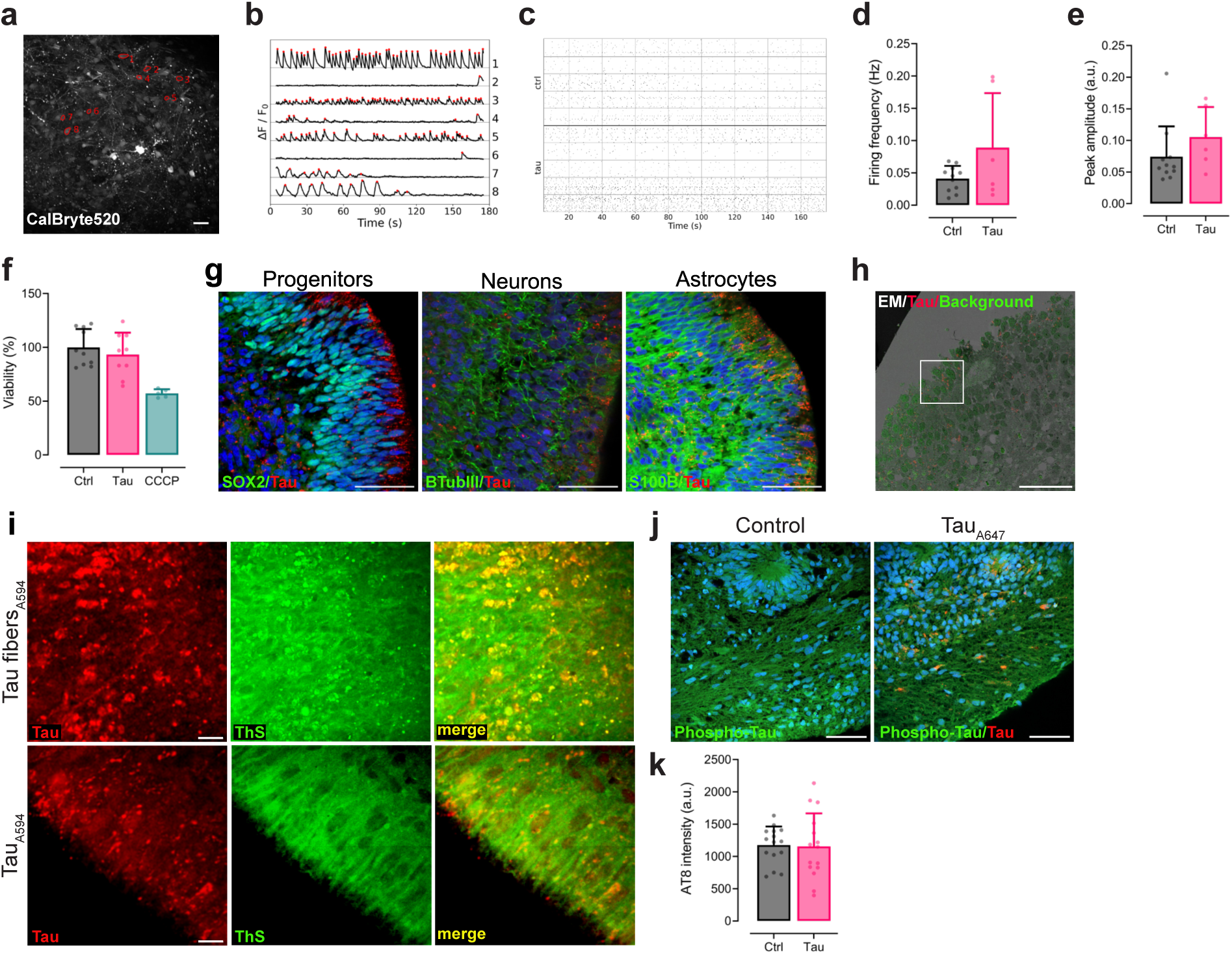
Functional validation, cell-type distribution, and absence of pathological tau species during early tau spreading in human forebrain organoids. (a) Representative calcium imaging recording of forebrain organoids using CalBryte520 as a calcium sensor (grayscale). Eight representatives segmented somata are highlighted in red. Scale bar, 50 µm. (b) Representative individual calcium traces corresponding to the segmented somas shown in (a). Red dots indicate detected peaks. (c) Raster plot showing segmented somas (each row) and detected peaks (each dot) over 180 s time lapse of all analyzed data sets for control (Ctrl, *n* = 11) and tau_A647_-treated organoids at 120h (tau, *n* = 6) calcium recordings. (d) Quantification of average firing frequency (Ctrl: 0.04 ± 0.01 Hz vs. Tau: 0.09 ± 0.08 Hz, *p* = 0.227) and (e) average peak amplitude (Ctrl: 0.07 ± 0.05 a.u. vs. Tau:0.11 ± 0.05 a.u., *p* = 0.149) in (c). Data were analyzed using the Welch’s test for (d) and Mann-Whitney test for (e). Data in (c-e) correspond to 6 individual organoids per condition, 3 experimental replicates. (f) Cell viability measured at 120 h following tau_A647_ conditioning. (Ctrl: 100 ± 17% vs. Tau: 93 ± 20%, *p* = 0.616). 24h 20µM CCCP treatment, which induces mitochondria dysfunction and subsequent cell death, reduced viability to 57% ± 4%. *n* = 10 (Ctrl), *n* = 9 (Tau) and *n* = 5 (CCCP) individual organoids, 4 experimental replicates. Data were analyzed by one-way ANOVA followed by a post-hoc Tukey’s test was used to determine P value. (g) Immunofluorescence staining showing internalized tau_A647_ (red) signal overlap with progenitors (SOX2⁺, green), neurons (βIII-tubulin⁺, green), and astrocytes (S100B⁺, green) markers. Nuclei in blue. Scale bars, 50 µm (h) Overview of correlative fluorescence and EM: in grey-scale, EM image; in red, fluorescent tau; in green, fluorescent background signal Boxed area highlights the ROI that is shown at higher magnification in Figure 1e. Scale bar, 50 µm. (i) ThS staining of organoid sections comparing brain organoids conditioned with tau fibers labeled with Alexa594 (tau fibers_A594_) and monomeric tau (tau_A594_) at 120h. Asterisks highlight ThS-positive tau fibers. (j) Immunostaining for phosphorylated tau (green) using AT8 antibody in control and tau_A647_ (red)-conditioned organoids at 120h. (k) Quantification of AT8 antibody fluorescence intensity in (j) (Ctrl: 1175 ± 287 a.u vs. Tau: 1154 ± 514 a.u., *p*=0.8904). *n* = 15 FOV per condition, 2 experimental replicates. P value was determined by an unpaired t test.

**Figure S2.**
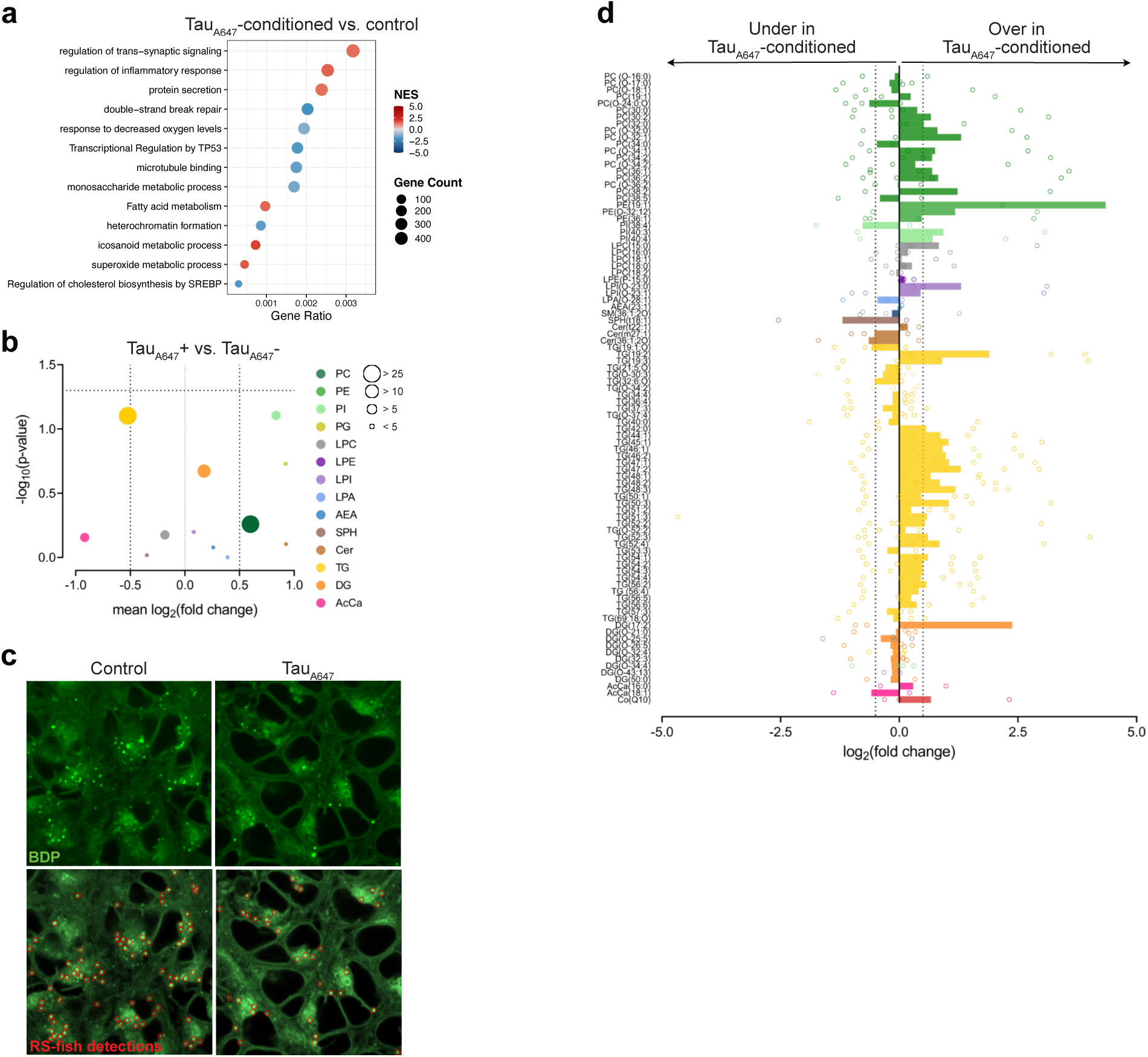
Global transcriptomic and lipidomic alternations upon tau spreading. (a) GSEA of transcriptomics data for tau-conditioned organoids compared to control, data represents *n* = 3 technical replicates. (b) Bubble plot presenting average log₂ fold change (tau_A647_+ vs tau_A647_-) for color-coded lipid classes (mean log₂ fold change versus -log₁₀ adjusted p value). Bubble size reflects number of lipid spices representing each lipid class. (c) Top: ROI from Fig. 2e comparing BDP staining in control and tau-conditioned organoids. Bottom: red circles highlight the RS-FISH detected spots. (d) Species-resolved log₂ fold change of lipid intensities for all detected lipids comparing tau-conditioned versus control, grouped and color-coded by lipid class. For (b) and (d), data represent independent organoid batches, *n* = 4 experimental replicates.

**Figure S3.**
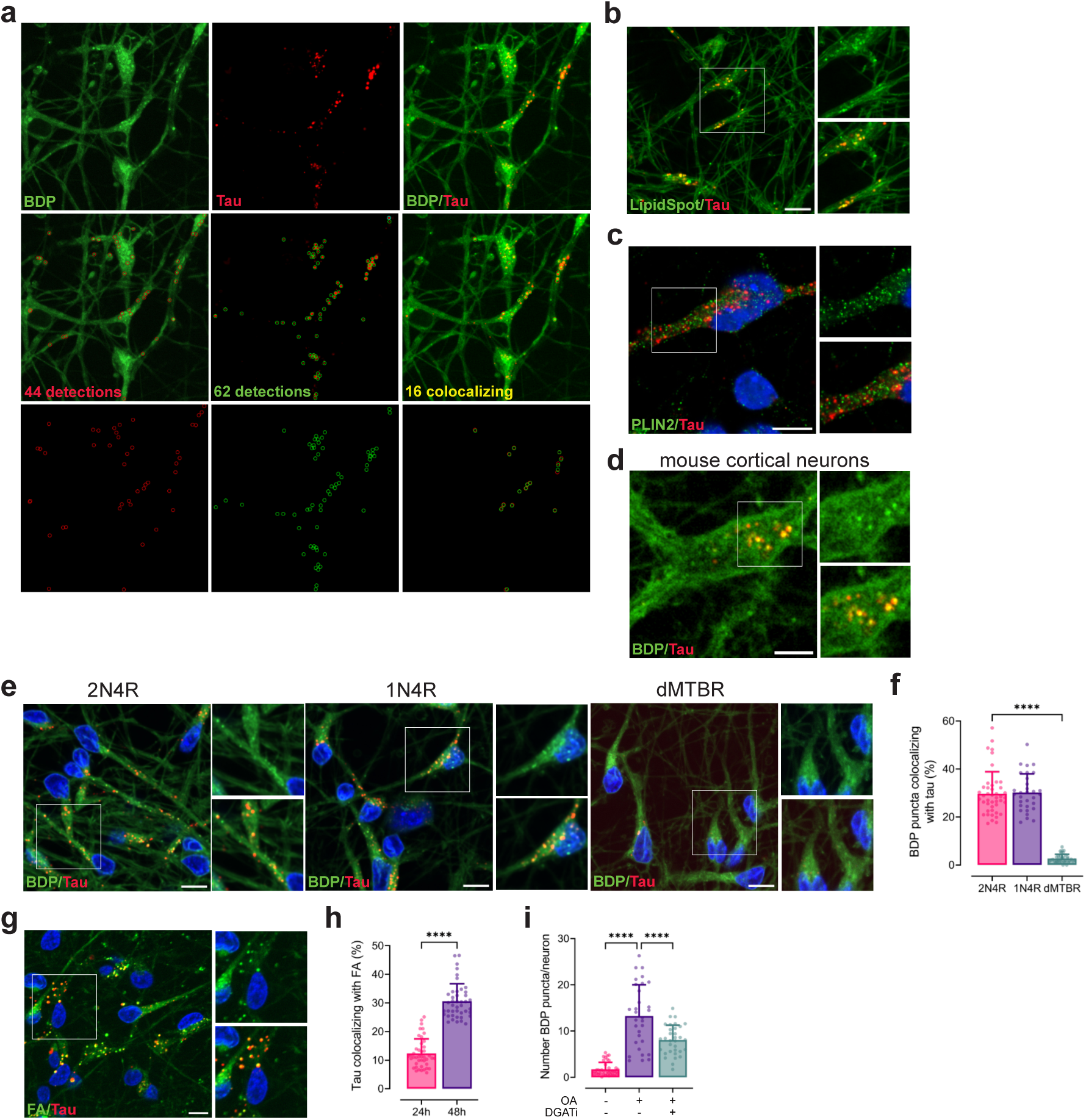
Spreading tau-neutral lipid interactions and endo-lysosomal trafficking. (a) Examples of RS-FISH puncta detection accuracy of live-cell imaging iPSC-derived forebrain neurons exposed to tau_A647_ (red) and stained with BDP (green): Top: raw images: center: overlap raw images with RS-FISH detections. Detected BDP puncta are shown as red circles and detected tau puncta as green circles. On the right side, only colocalizing puncta (centers closer than 0.5 µm) are shown. Bottom: for clarity, images whereonly detections are shown on a black background. (b) iPSC-derived forebrain neurons exposed to tau_A647_ (red) and stained with LipidSpot for neutral lipids (green). Insets highlight intracellular tau puncta colocalizing with neutral lipid–positive structures, 24.0 ± 11.6 % of tau puncta colocalize with LipidSpot puncta (*n* = 27 FOV, 3 experimental replicates). (c) Immunostaining of tau_A647_ (red)-conditioned neurons labeled for the LD-coating protein PLIN2 (green, left). Insets show partial colocalization: PLIN: 15.0 ± 4.9 % of tau puncta colocalize with PLIN puncta (coordinate-based colocalization, *n* = 27 FOV, 3 experimental replicates (d) Mouse hippocampal neurons conditioned with tau_A647_ (red) and labelled with BDP (green). Insets show higher-magnification views of colocalizing puncta, 24.1 ± 11.0 % of tau puncta colocalizing with BDP puncta (*n* = 53 primary neurons, 2 experimental replicates). (e) Representative images of neurons conditioned to different tau constructs (red) and stained with BDP (green): full-length 2N4R, 1N4R, and a C-terminal truncated tau variant lacking the microtubule-binding region (dMTBR). Insets highlight tau-BDP colocalization for each construct. (f) Quantification of tau colocalization with BDP for the constructs in (i) (2N4R: 29.7 ± 9.1 % vs. 1N4R: 30.1 ± 7.8 %, *p* > 0.9999 and 2N4R: 29.7 ± 9.1 % vs. dMTBR: 2.6 ± 1.8 %, *p* < 0.0001, n = 44 (2N4R), n = 31 (1N4R) and n = 36 (dMTBR) FOVs, 3 experimental replicates. Data were analyzed by Kruskal-Wallis test followed by a post-hoc Dunn’s multiple comparison test to determine P value. (g) Representative images of neurons enriched with fluorescent FA (BDP-C12) at 48 h after tau exposure. (h) Quantification of tau colocalization with FA comparing 24 h (Fig. 3d-e) and 48 h displaying Mander’s coefficient (% of tau overlapping lipid signal) (24h: 12.3 ± 5.1 % vs. 48h: 30.6 ± 6.1 %, *p* < 0.0001, *n* = 43 FOVs for each condition, 3 experimental replicates). P value was determined by Mann-Whitney test) (i) Quantification of the number of BDP puncta per neuron under control conditions (- OA -DGATi), oleic acid treatment (+ OA - DGATi), or OA plus DGAT1/2 inhibition (+OA +DGATi). (-OA – DGATi: 1.7 ± 1.4 vs. + OA – DGATi: 13.3 ± 6.7 number of BDP puncta, p < 0.0001 and; + OA – DGATi: 13.3 ± 6.7 vs. + OA + DGATi: 8.1 ± 3.2 number of BDP puncta, p < 0.0001, n = 39 (- OA - DGATi), n = 31 (+ OA – DGATi) and n = 33 (+ OA + DGATi) FOVs, 3 experimental replicates. Data were analyzed by one-way ANOVA followed by a post-hoc Tukey’s test was used to determine P value. For all images, nuclei are blue. Scale bars, 10 µm.

**Movie 3.** Time-lapse recording of tau_A647_ puncta (red) dynamics in neurons used to calculate tau trajectories shown in Fig. 4c. BDP (green) signal was used to identify cell morphology.

**Movie 4.** Time-lapse recording of tau_A647_ puncta (red) dynamics in astrocytes used to calculate tau trajectories shown in Fig. 4c. BDP (green) signal was used to identify cell morphology.

**Movie 5.** Time-lapse imaging of neurons sequentially conditioned with tau_A594_ (green) and tau_A647_ (red) shown in Fig. SI4b.

**Movie 6.** Time-lapse imaging of astrocytes sequentially conditioned with tau_A594_ (green) and tau_A647_ (red) shown in Fig. SI4b.

**Movie 7.** Time-lapse measurement of tau_A647_ (red)-conditioned neurons stained for tubulin (gray-scale) shown in Fig. SI4f

**Movie 8.** EM tomogram of electron tomography shown in Fig. SI4i

**Figure S4.**
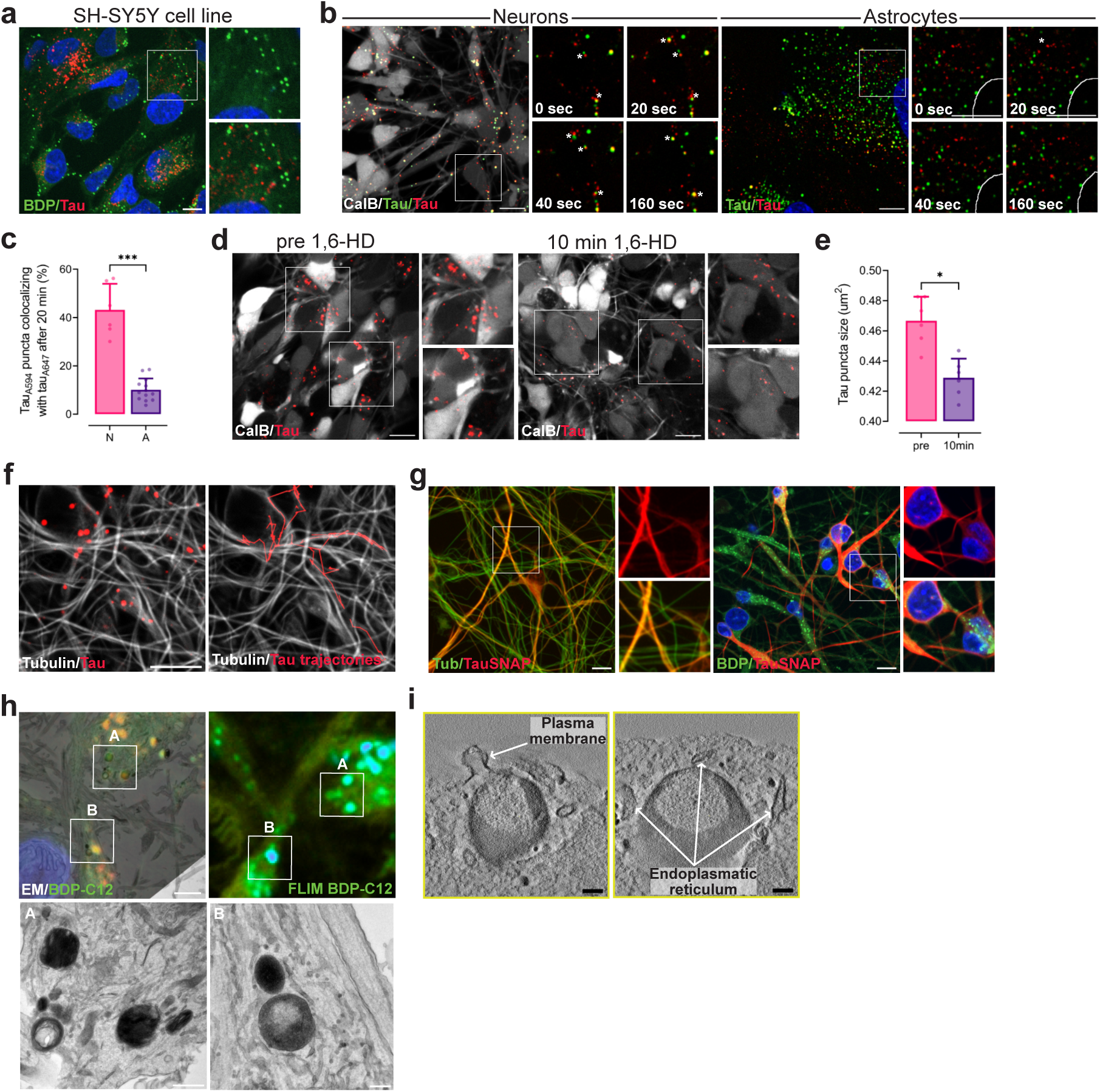
Spreading tau forms dynamic, lipid-associated assemblies distinct from intracellular tau. (a) SH-SY5Y neuroblastoma cell line exposed to tau_A647_ (red) and stained with BDP (green). Insets highlight not overlapping intracellular tau puncta and BDP-positive structures; 2.3 ± 1.1 % of tau puncta colocalize with BDP quantified by coordinate-based colocalization analysis. n = 26 FOVs, 3 experimental replicates. Scale bars, 10 µm. Nuclei are blue. (b) Representative frame of time-lapse imaging of neurons (left) and astrocytes (right) sequentially conditioned with tau_A594_ (green) and tau_A647_ (red). Insets highlight representative fusion and fission events, showing progressive merging of tau_A594_ and tau_A647_ signals over time in neurons. Time stamps indicate seconds after starting time-lapse imaging. Asterisks indicate representative fusion or fission events. Calbryte 520 (grey) or Hoechst 33342 (blue) were used to identify cell morphology. Scale bars, 10 µm. (c) Quantification of the percentage of tau assemblies double-positive for tau_A594_ and tau_A647_ after 20 min of imaging: N (neurons) 43.1 ± 10.9 % vs. A (astrocytes) 10.1 ± 4.6 % of tau, p = 0.0004, n = 6 (N) and n = 13 (A) time-lapse recordings of different FOVs, 3 experimental replicates. (d) Time-lapse imaging of neurons exposed to tau_A647_ (red) before (left) and 10 min after treatment with 1,6- Hexaendiol (1,6-HD). Insets highlight representative size tau puncta. Calbryte 520 (grey) was used to identify cell morphology. Scale bars, 10 µm. (e) Quantification of tau puncta size before and after 1,6-HD treatment (pre: 0.46 ± 0.02 µm^2^ vs. 10 min: 0.43 ± 0.01 µm^2^, *p* = 0.0313), *n* = 6 time-lapse recordings for each condition, 3 experimental replicates. Paired data was analyzed using Wilcoxon test. (f) Left: Representative frame of a time-lapse measurement of tau_A647_ (red)-conditioned neurons stained for tubulin (gray-scale)). Right: tau trajectories calculated by single-particle tracking (red) superposed to the tubulin signal (grey-scale), n = 14 time-lapse recording, 3 experimental replicates. Scale bar, 10 µm. (g) Left: overexpressed SNAP-tagged tau (red) iPSC-derived neurons stained for tubulin (green). Inset highlight a representative area were both fluorescence signals colocalize; 69.1 ± 5.2 % of tauSNAP signal overlaps tubulin signal analyzed using Mander’s coefficient intensity-based colocalization method, n = 32 FOVs, 3 experimental replicates. Right: Tau-SNAP (red) expressing neurons stained with BDP (green). Insets highlight a representative area where both fluorescence signals don’t overlap; 7.2 ± 4.7 % of the tauSNAP signal overlaps BDP signal analyzed by Mander’s coefficient. Scale bars, 10 µm. Nuclei are blue. (h) Top left: overview of FLIM-EM correlation of iPSC-derived neurons treated with BDP-C12 (green). Scale bar, 2 µm. Boxed areas (A and B) highlight the ROIs shown at higher magnification on the bottom row. Scale bars, (A) 500 nm and (B) 200 nm. Top right: corresponding FLIM image of BDP-C12. Color-scale bar indicates the range of lifetimes (ns). (i) Electron tomography of ROI C shown in Fig. 4g. The tomogram highlights contacts between tau-lipid assemblies and the plasma membrane (left) and the endoplasmic reticulum (right) Scale bars, 200 nm.

**Figure S5.**
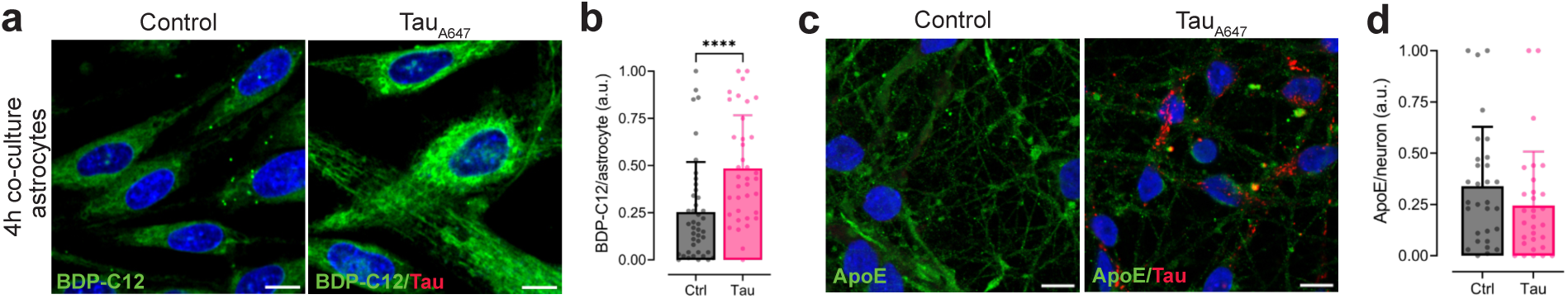
Characterization of tau-induced lipid transfer and ApoE regulation. (a) Representative images of neuronal BDP-C12 signal (green) in astrocytes co-cultured for 4 h with control or tau_A647_-conditioned neurons. (b) Quantification of neuronal BDP-C12 signal (green) in astrocytes after 4h of co-culture: Ctrl: 0.25 ± 0.27 vs. Tau: 0.48 ± 0.28 (a.u.), *p* < 0.0001; *n* = 42 (Ctrl) and *n* = 37 (Tau) FOVs, 3 experimental replicates Statistical analysis was done using Mann-Whitney test(c) (c) Immunofluorescence against ApoE (green) in neurons exposed to spreading tau (red) compared to control. (d) Quantification of ApoE fluorescence intensity per neuron: Ctrl: 0.46 ± 0.31 vs. Tau: 0.36 ± 0.31 (a.u.), *p* = ns; *n* = 27 (Ctrl) and *n* = 28 (Tau) FOVs, 4 experimental replicates Statistical analysis was done using a two-tailed unpaired t-test. Nuclei are in blue. Scale bars, 10 µm

**Figure S6.**
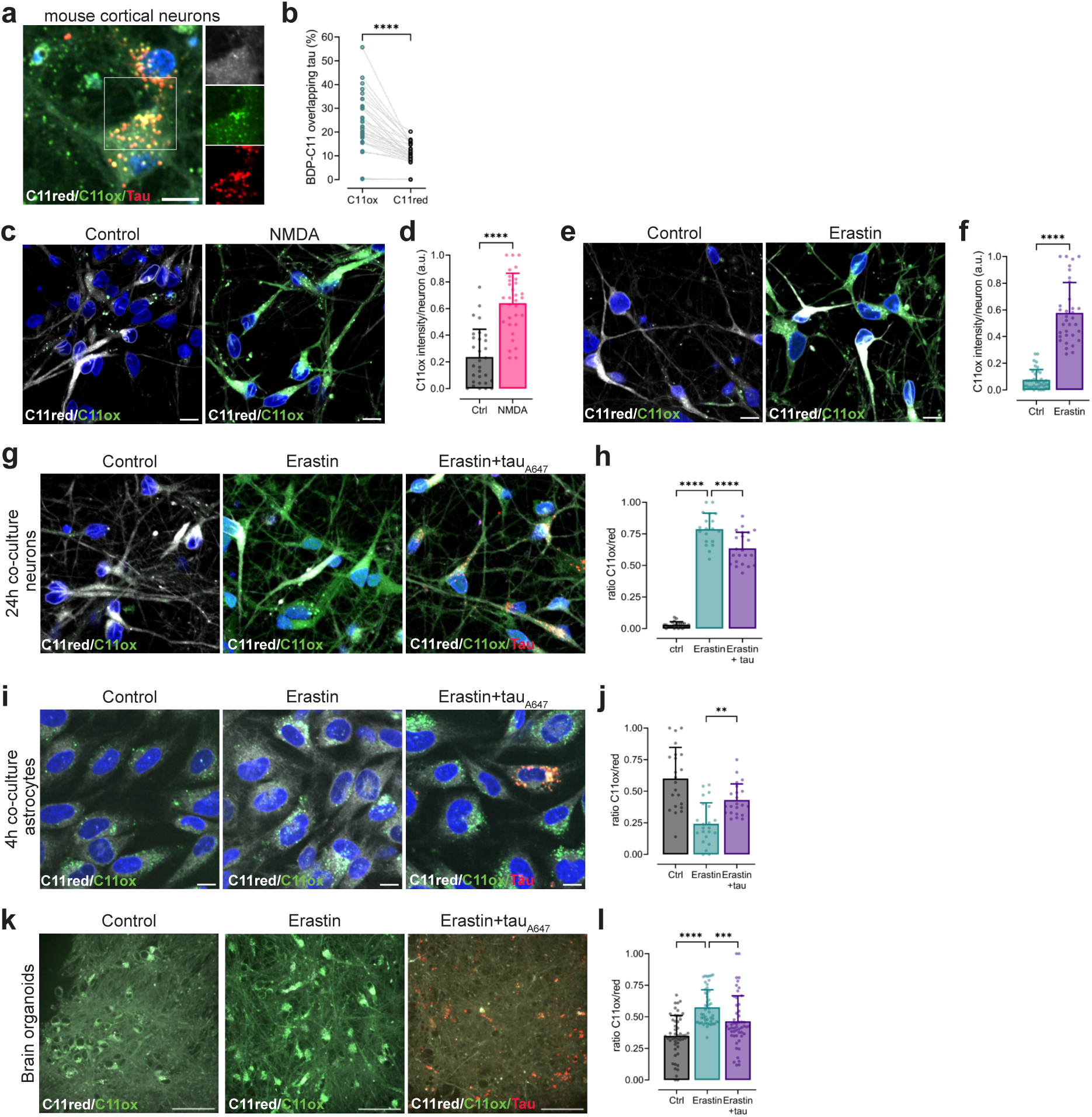
Tau-mediated handling of oxidized lipids under ferroptotic stress. (a) Representative images showing colocalization of tau (red) with BDP-C11(C11red (grey); C11ox (green) in primary mouse cortical neurons; insets highlight C11ox-tau puncta. (b) Percentage of tau signal verlapping with C11ox (17 ± 14 %,) versus C11red (8 ± 6 %; *p* < 0.0001) assessed by Mander’s coefficient; *n* = 46 FOVs, 2 independent experiments. Wilcoxon test was used to asses significance (c) Representative images of neurons supplemented with BDP-C11 (C11red, grey and C11ox in green) treated or untreated with NMDA. (d) Quantification of neuronal C11ox intensity in (c); Ctrl: 0.23 ± 0.21 a.u. *n* = 30 and NDMA: 0.64 ± 0.22 a.u. *n* = 31. N represent individual FOVs from 3 experimental replicates. P value (p < 0.0001) was calculated using a two-tailed unpaired t test. (e) Representative images of neurons supplemented with BDP-C11 (C11red, grey and C11ox in green) treated or untreated with erastin. (f) Quantification of neuronal C11ox intensity in (e); Ctrl: 0.08 ± 0.08 a.u. *n* = 37 and erastin: 0.58 ± 0.23 a.u. *n* = 34. N represent individual FOVs from 3 experimental replicates. P value (p < 0.0001) was calculated using Mann-Whitney test. (g) Representative imaging of BDP-C11-labeled neurons (C11red, grey and C11ox in green) co-cultured with astrocytes for 24 h under control conditions, after erastin treatment, or erastin treatment in the presence of spreading tau (red). (h) Quantification of neuronal C11ox/C11red ratio in (g): Ctrl: 0.03 ± 0.02 a.u. *n* = 23, Erastin: 0.79 ± 0.13 a.u. *n* = 19 and Erastin+tau: 0.63 ± 0.13 a.u. *n* = 22. Ns represent individual FOVs from 2 experimental replicates. P value (*p* < 0.0001) was determined by one-way ANOVA followed by a Tukey’s multiple comparison test: Ctrl vs. Erastin and Erastin vs. Erastin+tau, for both *p* < 0.0001. (i) Astrocytes co-cultured for 24 h with control neurons, erastin-treated neurons, or erastin-treated neurons exposed to spreading tau (red), labeled with BDP-C11(C11red (grey); C11ox (green)). (j) Quantification of astrocytic C11ox/C11red ratio comparing experimental conditions in (i): Ctrl: 0.60 ± 0.24 a.u., Erastin: 024 ± 0.16 a.u. and Erastin+tau: 0.43 ± 0.13 a.u. N = 22 individual FOVs for each condition from 2 experimental replicates. P value (*p* < 0.0001) was determined by one-way ANOVA followed by a Tukey’s multiple comparison test: Ctrl vs. Erastin (*p* < 0.0001) and Erastin vs. Erastin+tau (*p* = 0.00419). For images in (a, c, e, g and i) scale bar, 10 µm and nuclei are in blue. (k) Live-cell imaging of organoids treated with control, erastin, or erastin+tau and labeled with BDP-C11 (C11red (grey); C11ox (green)). Scale bar, 20 µm. (l) Quantification of organoid C11ox/C11red ratio comparing experimental conditions in (k); Ctrl: 0.35 ± 0.16 a.u. *n* = 52, Erastin: 0.58 ± 0.14 a.u. *n* = 50 and Erastin+tau: 0.46 ± 0.20 *n* = 52. Ns represent FOVs from 5 individual brain organoids per condition and 3 independent experiments. Statistical significance was determined using Kruskal-Wallis test (*p* < 0.0001) followed by post-hoc Dunn’s test; Ctrl vs Erastin (p < 0.0001) and Erastin vs Erastin+tau (p = 0.0005)

## Materials & Methods

### Cell culture of hiPSC lines

Human iPSCs (hiPSC) BIHi005-A was obtained from the MDC Pluripotent Stem Cell platform. The cells were cultured in standard hypoxic conditions (37 °C, 5% CO_2_, 5% O_2_ and 100% humidity), in Essential E8 Flex medium (Thermo Fisher Scientific, A2858501). Cultures were routinely monitored against mycoplasma contamination using PCR. Chemical passage of hiPSCs was performed at 70-80% confluency (every 3–4 days) in order to maintain log phase of growth and avoid induction of differentiation. Cells were detached with 0.5 mM EDTA (Thermo Fisher Scientific, 15575-020) in PBS (Thermo Fisher Scientific, 14190169) and seeded on Geltrex- (Thermo Fisher Scientific, A1413302) coated 6-well plates (Sarstedt, 833920).

### Generation and maintenance of neural stem cells (NSCs)

Neural stem cells (NSCs) were generated from the BIH005-A hiPSCs using a monolayer-based neural induction protocol employing dual SMAD inhibition^109^ iPSCs were dissociated into single cells with Accutase (Merck, A6964-100ML) and cells were seeded at a density of 5 × 10^4^ cells/cm² as a monolayer into geltrex-coated 6-well plate on E8 Flex media supplemented with 10 μM Rho-associated protein kinase (ROCK) inhibitor (Y-27632) (Hölzel Diagnostika, SEL-S1049-10MG) to prevent dissociation-induced apoptosis. When the culture reached 80-90 % confluency, the differentiation to NSCs was initiated by changing the media to Neural induction media (DMEM/F-12 (Thermo Fisher Scientific, 11330057) supplemented with N2 (Thermo Fisher Scientific, 17502048), B-27 (Thermo Fisher Scientific, 17504044), 100 U/mL penicillin and streptomycin (Thermo Fisher Scientific, 15140122), 10 μM SB431542 (VWR, 616461-5MG), a TGF-β signaling inhibitor, and 2 μM dorsomorphin (Enzo Life Sciences, ENZ-CHM141-0005), a BMP inhibitor). Medium was changed daily for 7 days, leading to the formation of a sheet of dense cells and the generation of NSCs. Next, cells were dissociated with Accutase into single cells and seeded geltrex-coated 6-well plates at a density of 150,000 cells/cm² in neural expansion media (NEM) (1:1 Advanced DMEM/F-12 (Thermo Fisher Scientific, 2634010):Neurobasal media with neural induction supplement (Thermo Fisher Scientific, A1647801) 100 U/mL penicillin and streptomycin and 5μM ROCK inhibitor). NPCs were maintained under proliferative conditions at 37 °C, 5% CO_2_ and 1% O_2_ in NEM media and received daily media changes. Cells were passaged at 70%-80% confluence using Accutase and seeded at 1.5 × 10^5^ cells/cm² in the presence of 5 μM ROCK inhibitor. NSCs were routinely monitored for characteristic morphology, rosette formation, growth characteristics and used for experiments within 9 passages. NSC identity was confirmed by expression of PAX6 (BioLegend, 901301) and SOX2 (Merck, AB5603), canonical neural stem cell markers, by flow cytometry and immunostainings. For NSC banking, cells were cryopreserved at early passages (passage 2 to 4). NSCs were dissociated into single cells using Accutase and cryopreserved in Bambanker (Nippon Europe, BB05) freezing medium containing 10 μM ROCK inhibitor and frozen using a controlled-rate cooling procedure before storage in liquid nitrogen.

### Tau-SNAP line generation

To generate tauSNAP PiggyBac hiPSC line (Fig. SI4), the BIHi005-A hiPSC parental line was transfected with tauSNAP Piggybac and transposase plasmid using Lipofectamine Stem (Thermo Fisher Scientific, STEM00008) One day prior to transfection, BIHi005-A hiPSCs colonies, at 70-80% confluency, were dissociated into single cells with Accutase. Next, 5 × 10^4^ cells were seeded per well onto Geltrex-coated 6- well plates in E8 Flex media supplemented with 10 μM ROCK inhibitor and 0.5x CloneR2 (STEMCELL Technologies, 100-0691) to promote hiPSC survival and clonal expansion. After 16 h, the media was replaced with E8 Flex supplemented with 0.5x CloneR2 and without ROCK inhibitor. Transfection was performed as follows: for each sample, 144 μl Opti-MEM (Thermo Fisher Scientific, 31985062) was mixed with 6 μl Lipofectamine STEM. In parallel, 1,000 ng tauSNAP PiggyBac plasmid and 400 ng transposase plasmid (1,400 ng total DNA) were diluted in 144 μl Opti-MEM. The diluted Lipofectamine and DNA solutions were combined at a 1:1 ratio and incubated for 10 mins at room temperature. A total of 300 μL of the mixture was then added dropwise to the hiPSC cultures. 20 h post-transfection, the media was replaced with E8 Flex supplemented with 0.5x CloneR2. Transfected tauSNAP PiggyBac hiPSCs were subsequently selected for one week in E8 Flex media containing 0.7 μg/ml puromycin (Thermo Fisher Scientific, A1113803).

### Forebrain neuronal differentiation from NSCs

Forebrain neuronal differentiation was performed using a commercially available STEMdiff forebrain neuron differentiation kit (STEMCELL Technologies, 08600) according to manufacturer’s instructions. Differentiation was initiated from NSCs at passage 6 to 8, either directly following neural induction or after recovery from cryopreservation. At day *in vitro* (DIV) 0, to generate neuron precursors, cells were seeded into poly-L-ornithin (Merck, P4957 - 50 mL)/laminin (Merck, L2020-1MG)-coated 6-well plate at a density of 1.2 × 10^5^ cell/cm^2^ and allowed to equilibrate under NSC maintenance conditions (NEM media and hypoxia) prior to initiation of forebrain differentiation. Next day, differentiation was initiated by replacing NEM for STEMdiff Forebrain Neurons Differentiation medium and moved to normoxia conditions, 37°C and 5% CO_2_. Cells were cultured for 6 days with daily media changes. At DIV6, forebrain precursors were dissociated with Accutase and seeded at 8 × 10^4^ cell/cm^2^ in STEMdiff Forebrain Neuron Maturation Medium (STEMCELL Technologies, 08605) on poly-L-ornithin/laminin-coated vessels. For live-cell imaging experiments and immunostainings, neurons were seeded in 8-well imaging dishes (ibiTreat, ibidi, 80806), for co-culture experiments in round glass coverslips (12mm diameter, 1.5mm thickness (VWR, 112520)) treated with nitric acid (Merck, 1004410250) and for correlated electron microscopy in 35mm glass bottom grids (ibidi, 81148). Media was changed every 2-3 days. Neurons were considered mature and used for experiments after DIV20. Neuronal maturity was assessed by calcium imaging and immunostainings of canonical neuronal markers (MAP-2 (Synaptic Systems 188002) and βIII-tubulin (BioLegend, 801201).

### Astrocyte differentiation from NSCs

Astrocyte differentiation was performed using a protocol adapted from Alisch et al^110,111^. Differentiation was initiated from NSCs at passage 4 directly following neural induction and avoiding intermediate cryopreservation step, which can bias cell survival and compromise lineage commitment. NSCs were dissociated into single cells with Accutase and seeded in a geltrex-coated 6-well plate at 5 × 10^4^ cell/cm^2^ in NEM medium in the presence of 5 µM ROCK inhibitor and kept in hypoxic conditions (37 °C, 5% CO_2_, 5% O_2_ and 100% humidity). Next day, NEM was replaced by Astrocyte differentiation media (DMEM/F-12, 2mM GlutaMax supplement (Thermo Fisher Scientific, 35050061), 10% fetal bovine serum (FBS) (Thermo Fisher Scientific, A5256701), N2 supplements, 20 ng/ml CNTF (Thermo Fisher Scientific, 450-13-50UG) and 100 U/ml penicillin and streptomycin) and cells were moved to an incubator with normoxia conditions and 5% CO_2_. Media was changed every 2-3 days. Differentiating astrocytes were passaged at 70%-80% confluence using Accutase. During the first 1 to 3 weeks of differentiation, cells were highly proliferative and subsequently become post-mitotic. At DIV40, astrocytes were considered mature. The expression of canonical astrocytic markers (GFAP (Agilent, Z033429-2), S100β (Abcam, ab52642) and the absence of Nestin (Proteintech, 19483-1-AP)) was checked by immunostaining and cells were used for downstream analysis.

### Derivation of cortical brain organoids

Brain organoids were generated at the BIMSB MDC Organoid platform from two different hiPSC lines: the XM001^112^. and Gibco Human Episomal iPS line 1E6 (Thermo Fisher Scientific. A18944, lot no. 2036936) using the forebrain protocol following a modified version of Walsch et.al.^113^. iPSCs were cultured in standard hypoxic conditions (37 °C, 4% CO_2_, 4% O_2_ and 100% humidity), in E8 Flex medium. On DIV0, cells were dissociated into a single-cell suspension using TrypLE (Thermo Fisher Scientific, 12604013), 9 × 10^3^ cells were seeded per well in 96-well plates with 100 µl of Essential 8 (E8) medium (Thermo Fisher scientific, A1517001) containing 50 µM ROCK inhibitor (Tocris, 1254) and 5 µM XAV939 (Enzo Life Sciences, BML- WN100-0005). Cells were kept in a hypoxia condition. On the following day, 80% of the medium was replaced with Essential 6 (E6) medium (Thermo Fisher Scientific, A1516401) supplemented with 0.1 µM LDN (Tocris, 6053), 5 µM XAV, and 10 µM SB (STEMCELL Technologies, 72234). On day 5, the medium was replaced with E6 medium supplemented only supplemented with 5 µM XAV939, 0.1 µM LDN-193189, and 10 µM SB431542. Embryoid bodies (EBs) were then transferred to normoxic conditions. Between DIV7 and 9, liquid embedding of the organoids was performed using 2% Matrigel (Corning, 356234) diluted in organoid differentiation medium. This differentiation medium consisted of a 1:1 mixture of DMEM/F12 (Thermo Fisher Scientific, 11320074) and Neurobasal medium (Thermo Fisher Scientific, A3582901), supplemented with N2 supplement (STEMCELL Technologies, 07152), B27 supplement minus vitamin A (STEMCELL Technologies, 05731), 3.2 µM insulin (Sigma-Aldrich, I9278), 0.05 µM 2-mercaptoethanol (Thermo Fisher Scientific, 35050038), 1× GlutaMAX (Thermo Fisher Scientific), MEM non-essential amino acids (Sigma-Aldrich, M7145), and leukemia inhibitory factor (LIF; PeproTech, 300-05-25UG). Embedded organoids were cultured in ultra-low-attachment 6-well plates on an orbital shaker (80 rpm) until day DIV15. From DIV15 onward, the medium was replaced with organoid maturation medium, consisting of a 1:1 mixture of DMEM/F12 and Neurobasal supplemented with N2, B27 supplement with vitamin A (STEMCELL Technologies, 05711), 3.2 µM insulin, 0.05 µM 2-mercaptoethanol, GlutaMAX, MEM non-essential amino acids, 0.2 mM ascorbic acid (Sigma-Aldrich, A4544), 20 ng/mL brain-derived neurotrophic factor (BDNF; STEMCELL Technologies, 78005.3), 20 ng/mL glial cell line-derived neurotrophic factor (GDNF; STEMCELL Technologies, 78058.3), and cyclic AMP (cAMP; Tocris, CAS 16980-89-5). Brain organoid experiments were performed using independent differentiation batches derived from the two hiPSC lines (XMOO1 and Gibco Human Episomal iPS line 1E6) to unsure reproducibility. Functional validation of successful cortical organoid derivation was performed at DIV60 by calcium imaging (Fig. SI1a-e) and upon passing quality control organoids were used for further downstream experiments.

### Cell culture of immortalized cell line

CCF-STTG1 cell line, an immortalized human astrocytoma line, was obtained from ATCC. Cells were grown in RPMI 1640 ATCC medium (Thermo Fisher Scientific, A1049101) supplemented with 10% FBS and 100 U/mL penicillin and streptomycin. SH-SY5Y cells were cultured in DMEM plus 10% FBS, 100 U/ml penicillin, and 100 μg/ml streptomycin. Both cell lines were grown at 37°C under a humidified atmosphere of 5% CO_2_. For maintenance, cells were passaged upon reaching ∼ 95% confluency, once or twice a week, with 0.05% Trypsin-EDTA (Thermo Fisher Scientific, 25300054) and seeded in 10 cm cell culture treated dishes (Sarstedt, 833902). CCF-STTG1 used in experiments were seeded at a density of 5 × 10^4^ cells/cm^2^ in geltrex-coated glass-bottom 8-well ibidi chambers or 24-well plates (ibidi, 82427).

### Primary cortical neurons

Primary cortical neurons were maintained in co-culture with feeder astroglia. Both were prepared from C57BL/6N mouse pups as previously described in Kaech and Banker^114^. In brief, cortices were isolated from wildtype C57BL/6N mice (P0 for neurons and P2 for astroglia). For astroglia, tissue was dissociated in 0.5% Trypsin solution (Life Technologies, 11580626) and then seeded in 12-well plates (∼30.000 cells per well), until confluency. For neurons, tissue was dissociated in papain (Worthington, LS003126) and the single cell suspension seeded in a previously Poly-L-Lysin (Sigma Aldrich, P2636)-coated glass coverslip. The neuronal coverslips were then placed on top of a monolayer of feeder astroglia, using paraffin dots to prevent direct contact of both cellular layers. The co-culture was maintained for to 14 days in vitro in Neurobasal-A (Life Technologies, 10888022) medium supplemented with 2% B-27 (Life Technologies, 17504044), 100U/ml pen/strep and 0.5 mM GlutaMAX-I (Life Technologies, 35050061) at 37°C and 5% CO_2_.

### Plasmids

pET-HT 2N4R and 1N4R tau plasmids for bacterial expression were a kind gift from Elizabeth Rhoades and were used for tau protein purification. For site-specific fluorescence labeling, the native cysteines (Cys291 and Cys322) were mutated to serines, enabling the introduction of a single cysteine residue at the desired position. Depending on the experimental design, either tauT17C or tauS433C was used. The suitability of these labeling sites has been validated previously (REF). pET-HT tau deltaMTBR (microtubule binding region) was generated by site-directed mutagenesis (Q5® Site-Directed Mutagenesis Kit, New England Biolabs, E0554S) of the full-length tau 2N4R construct. A premature STOP codon was introduced at the position 245, resulting in the expression of a truncated protein lacking MTBR domain and the C-terminus. To generate pcDNA3.1 tau-SNAP for mammalian expression, the open reading frame (ORF) of the 2N4R tau plasmid and the ORF of the SNAP tag from the pcDNA3.1 Halo-SNAP-mGluR ^114^ plasmid were amplified by PCR and assembled into a pcDNA3.1(+) backbone by Gibson Assembly (NEBuilder HiFi DNA Assembly Master Mix, New England Biolabs, E2621L) according to the manufacturer’s instructions. ePB_Puro-TT- RFP Tau-SNAP PiggyBac plasmid was generated by cloning the ORF of tau-SNAP (from pcDNA3.1 tau- SNAP plasmid) into the ePB_Puro-TT-RFP PiggyBac backbone using Gibson Assembly. Primers containing 5′ homology arms to generate overlapping ends were designed using NEBuilder Assembly Tool (New England Biolabs) and PCR fragments were amplified using Q5 High-Fidelity DNA Polymerase (New England Biolabs, M0491L). All mutations were confirmed by Sanger sequencing (Eurofins Genomic) and newly generated plasmids verified by whole-plasmid Nanopore sequencing (Plasmidsaurus).

### Tau purification and labeling

Tau isoform, 2N4R and 1N4R, and domain variants, delta MTBR, were expressed in E. coli BL21 cells from pET-HT 2N4R, 1N4R and deltaMTBR tau plasmids. The parent plasmid includes an N-terminal His tag with a tobacco etch virus (TEV) protease cleavage site for purification. For all constructs protein expression was induced with 1mM IPTG (Biomol, I8500.10) at OD 0.4 overnight at 16°C. Purifications was based on methods reported previously ^115,116^. Briefly, cells were lysed in presence of 1 mg/ml lysozyme (Merck, L6876- 1G), EDTA-free protease inhibitor (Merck, 11836170001) and 1 mM PMSF (Merck, 10837091001) by sonication, and the cell debris was pelleted and removed by centrifugation. The supernatant was incubated with nickel-nitrilotriacetic acid resin (Merck, 70666-4), and the recombination protein was eluted with 500 mm imidazole (Merck, 8.142.23). The His tag was removed by overnight incubation at 4°C with TEV proteinase (purchased from inhouse protein purification facility). Uncleaved protein was removed by a second pass over the nickel-nitrilotriacetic acid column. Remaining contaminants were removed using size exclusion chromatography on a Superdex200 Increase 10/300 GL (Merck).

For site-specific fluorescence labeling, freshly purified tau (typically 300-500 μl of ∼100 μM protein) was incubated with 1 mM dithiothreitol (DTT) (Thermo Fisher Scientific, 15508013) for 30 min at room temperature to reduce the cysteines. The protein solution was buffer exchanged into 20 mM Tris (pH 7.4), 50 mM NaCl, and 6 M guanidine hydrochloride (GdmCl) (Merck, G3272) and concentrated to 1 mL using Amicon Ultra concentrators (10,000 Da MWCO) (Merck, UFC9010). The protein was incubated with 5 × molar excess Atto700 (Leica, AD-700-41), Alexa647 (Thermo Fisher Scientific, A20347) or Alexa594 maleimide (Thermo Fisher Scientific, A10256)) overnight at 4°C under stirring conditions. The labeled protein was concentrated and buffer exchanged into 20 mM Tris pH 7.4, 50 mM NaCl using an Amicon Ultra concentrator (10,000 Da MWCO), with final removal of unreacted dye and remaining GdmCl by passing the solution through two coupled HiTrap desalting columns (Merck, 17-1408-01) equilibrated with 20 mM Tris pH 7.4, 50 mM NaCl. The fractions containing labeled tau were pooled, diluted to 30 μM in the same buffer, snap-frozen and stored at -80 °C. Tau 2N4R was labelled at either N(TauT17C) or C-terminus (tauS433C) cysteines and both were used in subsequent experiments. Tau 1N4R isoform was labeled at the C-terminus (tauS433C) and tau deltaMTBR at the N-terminus (tauT17C). Tau_A700_, tau_A647_ and tau_A594_ were use according to the available and matched laser wavelength for the specific microscopic set ups used and to minimize spectral overlap with other fluorophores. Final protein concentration was measured by single molecule fluorescence correlation spectroscopy (FCS) (PicoQuant MicroTime 200) providing a quantitative assessment on the protein concentrations used in the study. Human tau pre-formed fibers (Acrobiosystems, TAU-H5115) stored at -80°C were let to temperate to room temperature and were labelled with 150 µM AlexaFluor594 NHS Ester ((Thermo Fisher Scientific, A37572) for 1h. The labelling reaction was stopped by adding 100 mM Tris-HCl pH 7.5. After 2 washing steps with PBS by centrifugation (10 minutes at 15.000g), fibers were resuspended in PBS and sonicated for 30 min at 37°C in a water bath, aliquoted and stored at -80°C until further use.

### Reconstitution of lipid probes and fluorescent lipid analogs

BODIPY 493/503 (Thermo Fisher Scientific, D3922), BODIPY 581/591 C11(Thermo Fisher Scientific, D3861) were dissolved in DMSO (Thermo Fisher Scientific, D12345) at a concentration of 8 mM. N-(4,4- Difluoro-5,7-Dimethyl-4-Bora-3a,4a-Diaza-s-Indacene-3-Dodecanoyl) SphingBosyl Phosphocholine (BODIPY FL C12-Sphingomyelin, Thermo Fisher Scientific, D7711) was supplied at 1mg/ml solution in DMSO. 4,4-Difluoro-5-(2-Thienyl)-4-Bora-3a,4a-Diaza-s-Indacene-3-Dodecanoic Acid (BODIPY 558/569 C12, Thermo Fisher Scientific, D3835) was dissolved at 2 mM in DMSO. All lipid probes and fluorescent lipid analogs described until now were aliquoted and stored at -20°C, protected from light, following manufacturers recommendation. Oleic acid (OA, Merck, O1008) was dissolved in 50% ethanol at a 100μM concentration and kept at 4°C. 23-(dipyrrometheneboron difluoride)-24-norcholesterol (TopFluor Cholesterol, Merck, 810255P) and CholEsteryl 4,4-Difluoro-5,7-Dimethyl-4-Bora-3a,4a-Diaza-s-Indacene- 3-Dodecanoate (CholEsteryl BODIPY FL C12, Thermo Fisher Scientific, C3927MP) were first dissolved in chloroform and the solvent was evaporated by blowing nitrogen followed by vacuum desiccation until a lipid film was formed at the bottom of the glass container. Before the experiment the lipid film was reconstituted into a lipid solution at 3.5mM and 1mM in DMSO for TopFluor Cholesterol and CholEsteryl BODIPY FL C12, respectively.

### Single-cell dissociation of cortical brain organoids

Brain organoids were conditioned with 20 nM tau_A647_ in brain organoid maturation medium or with 25 µg/ml Transferrin-A594 (Tf_AF594_) (Thermo Fisher Scientific, T13343). After 24h, media was replaced completely to remove any excess tau or transferrin that did not internalize. Organoids were then maintained in culture under standard conditions until the specified time points. Treatments such as Heparinase I and III, 100U/ml (Merck, H3917), and RAP, 500nM (Enzo Life Science, BML-SE552-0100), were added directly to the culture medium at each medium change, when appropriate. To proceed to flow cytometry, brain organoids were dissociated into single cells using the Neural Tissue Dissociation Kit (P) (Miltenyi Biotec, 130-092-628). Briefly, organoids were dissociated individually: first mechanically using a scalpel, followed by enzymatic digestion with Enzyme Mix 1 and incubation for 10 min at 37°C with gentle shaking. Samples were then further mechanically dissociated by pipetting and incubated for an additional 10 min at 37°C, with Enzyme Mix 2. Dissociation was terminated by adding DMEM/F12 supplemented with 10% FBS, and the cell suspension was filtered through a 40 µm cell strainer.

### Flow cytometry analysis and cell sorting

Single cell suspension was centrifuged at 300 x g for 5 min and resuspended in FACS buffer (1% BSA (Thermo Fisher Scientific, 10326640), 25 mM HEPES (Merck, H0887-100ML), 0.5 mM EDTA (Thermo Fisher Scientific, 15575020), 10 µM ROCK Inhibitor in DPBS). DAPI (Thermo Fisher Scientific, 62248) (0.2 µg/ml) was added to stain the nuclei of dead cells. Different instruments were used for flow cytometry analysis and cell sorting. To quantify tau spreading in brain organoids (data shown in Fig. 1f-g), flow cytometry was performed using using the LSRFortessa™ X-20 Cell Analyzer (BD Biosciences) equipped with the FACSDiva™ Software (v9.0.1). A total of 500,000 events per sample were acquired using the following settings: 405 nm excitation, 450/50 BP emission filter for DAPI; 561 nm excitation, 610/20 BP emission filter for Tf_A594_; and 635 nm excitation, 670/30 BP emission filter for tau_A647_. The FCS files analyzed in FlowJo (v10.10.0): forward scatter (FSC) and side scatter (SSC) were used to exclude debris and clumps, followed by singlet discrimination using FSC-A (area) versus FSC-H (height), and SSC-A versus SSC-H. Live cells (DAPI-negative), tau_A647_-positive (tau+) and Tf_A594_-positive (Tf+) population in each condition was quantified based on gates defined using unstained control samples. For sorting tau+ and tau- cell population for lipidomics and transciptomics (data shown in Fig.2 and Fig. SI2) single-cell suspensions were processed using a BD FACSAria™ Fusion Flow Cytometer (BD Biosciences) equipped with a 100 µm nozzle. The following settings were using: 405 nm excitation, 450/50 BP emission filter for DAPI and 635 nm excitation, 670/30 BP emission filter for tau_A647_. Cells were gated sequentially following the same strategy described above. Sorted cells were collected in FACS buffer at 4°C in 15 ml tubes and used immediately for downstream applications, including transcriptomics or lipidomics analysis.

### Lipidomic, lipid extraction, processing and analysis

For lipidomics analysis, at least 200,000 FACS-sorted cells per condition were collected. Sorted cells were centrifuged, and the resulting pellet was snap-frozen and stored at -80°C until lipid extraction. Lipids were extracted with the Folch method. Briefly, 600 μl of chloroform:methanol. (2:1, v/v), containing 7.5 μl of SPLASH Lipidomix standards (cat. # 330707, Avanti Research, USA) was added to each sample. Following sonication, 130 μl of water was added to induce phase separation. After centrifugation (5 min, 5,000 x g), 400 μl of the non-polar phase was collected. Subsequently, 5 μl of HCl (1M) and 400 μl of chloroform were added. After mixing and centrifugation (5 min, 5000 x g) to achieve phase separation, an additional 400 μl of non-polar phase was collected and combined with the initial extract, yielding 800 μl in total. The combined lipid phase was dried under nitrogen flow and stored at -80 C until sample processing. Prior to LC-MSMS analysis, dried lipids were resuspended in 50 μl of isopropanol:acetonitrile (1:1, v/v). Chromatographic separation was performed using a Vanquish Horizon UHPLC system (Thermo Fisher Scientific) equipped with an Accucore Vanquish C18+ reversed-phase column (2.1 x 100mm, 1.5 µm bead size Thermo Fisher Scientific, USA). A 2 µl injection volume was used for each run. Raw LC–MS data were processed using Compound Discoverer 3.4 (Thermo Fisher Scientific) with an adapted version of the untargeted lipidomics workflow. Compound annotation was performed hierarchically using the LipidBlast and LipidSearch spectral libraries. A custom R-script was used to perform post-processing and statistical analysis.

### Transcriptomics, RNA extract and transcriptomic analysis

For RNA extraction, 20,000 FACS-sorted cells were collected in FACS buffer containing RiboLOck RNase Inhibitor (Thermo Fisher Scientific, EO0381) for each condition. Total RNA was isolated using the TRIzol reagent. Briefly, single-cell suspension was homogenized by bead beating in 1 ml of TRIzol reagent (Thermo Fisher Scientific, 15596026) and incubated for 5 min at room temperature to allow complete dissociation of nucleoprotein complexes. Subsequently, 0.2 ml of chloroform (Carl Roth, 3313.2) per 1 ml of TRIzol was added, samples were shaken vigorously, incubated for 2–3 min at room temperature, and centrifuged at 12,000 × g for 15 mins at 4°C. The clear aqueous phase was carefully transferred to a new tube without disturbing the interphase, and 0.5 ml of isopropanol (Carl Roth, 0080.3) per 1 ml of TRIzol was added. Samples were mixed by inversion and incubated for 10 min at room temperature, followed by centrifugation at 12,000 × g for 30 min at 4°C to pellet the RNA. The supernatant was removed, and the RNA pellet was washed with 85% ethanol, followed by brief air-drying. RNA was dissolved in RNase-free water.

Bulk RNA-seq data analysis was performed in the R statistical computing environment (v4.5.2). Differential gene expression analysis was carried out using the DESeq2 package (v1.50.0). The raw count matrix was filtered to remove genes with counts <10, followed by normalization using the DESeq2 variance stabilizing transformation (VST) to reduce heteroscedasticity. Principal component analysis (PCA) was performed on the transformed data to assess sample clustering. Significantly differentially expressed genes (DEGs) were identified using a Benjamini–Hochberg false discovery rate (FDR)-adjusted p value <0.05 and an absolute log2 fold change >0.5. Log2 fold-change shrinkage was applied using the adaptive shrinkage (“ashr”) method implemented in the lfcShrink function to obtain stabilized effect size estimates. Gene identifiers were annotated using the biomaRt package (v2.66.0) by querying the Ensembl human gene dataset to retrieve HGNC gene symbols and Entrez Gene IDs. A database of lipid-related genes was compiled from lipid-associated gene sets obtained from the Molecular Signatures Database (MSigDB, v3.0). DEGs were intersected with this database, and their distribution was visualized using volcano plots. Gene set enrichment analysis (GSEA) was performed using the clusterProfiler package (v4.18.1). Redundancy among enriched Gene Ontology (GO), Reactome Pathway (RP), and KEGG terms was reduced based on semantic similarity and hierarchical relationships. Representative parent terms were selected to summarize related biological processes, and significantly enriched pathways were visualized using manually curated dot plots generated with ggplot2 (v4.0.0).

### Viability assay

Cell viability was determined using a resazurin reduction assay. Individual brain organoids were placed in a single well of a 24-well plate in brain organoid maturation media. After tau conditioning, resazurin (biomol, Cay14322-5) was added at a final concentration 0.01 µg/ml and organoids were incubated for 4h at 37°C. Fluorescence was measured using a microplate reader (Molecular Devices, SpectraMax iD5) (excitation 560 nm, emission 590 nm). As a positive control for reduced viability, organoids were treated with 20µM carbonyl cyanide m-chlorophenyl hydrazone (CCCP) (Merck, 215911-250MG) for 24h. Raw fluorescence values were corrected for background and normalized to the weight of each individual organoid to account for size differences. Data were subsequently expressed relative to untreated controls.

### Live-cell imaging

When appropriate, brain organoids or neuronal cultures were conditioned with 20 nM tau_A647_ for 16h or 24h and prior to imaging, live cell stainings (described below) were performed. Live-cell staining and live-cell imaging experiments were conducted in phenol red-free and serum-free BrainPhys Imaging Optimized Medium (STEMCELL Technologies, 05796) unless otherwise stated, in environmentally controlled conditions at a temperature of 37°C and 5% CO_2_. When indicated, nuclei were labeled with 2 µM Hoechst 33342 (Merck, B2261-25MG) for 15 min at 37°C prior to imaging. When possible, FOVs were selected using a channel that allowed blinded image acquisition. FOVs were acquired across the entire well to ensure representative sampling

#### Calcium imaging

For calcium imaging experiments, brain organoids were incubated with 5µM CalBryte520 AM (ATT Bioquest, ABD-20651) and 0.035% Pluronic acid (Thermo Fisher Scientific, P3000MP) for 1h at 37°C and 5% CO_2_ on an orbital shaker (80 rpm). Organoids were allowed to equilibrate to the imaging chamber and settle onto 35 mm glass-bottom dishes for 5 min prior to initiating calcium measurements. Similarly, calcium imaging of hiPSC-derived forebrain neuronal cultures was performed by incubating neurons with 5µM CalBryte520 AM and 0.035% Pluronic acid, for 30 mins. Time-lapse recordings were acquired in the CalBryte520 channel, and the average amplitude and firing frequency of calcium oscillations were quantified.

#### Neutral lipid imaging

Brain organoids treated with 20 nM tau_A647_ were incubated with 2 µM BODIPY 493/503 (BDP; Thermo Fisher Scientific, D3922) for 1h under standard culture conditions. Prior to imaging, organoids were washed three times with imaging medium. hiPSC-derived forebrain neuronal and astrocytic cultures, mouse primary neurons, CCF-STTG1 cells and SHSY5Y cells were stained using the same protocol with shorter a incubation time (30 min). As an alternative approach, neutral lipids were stained using LipidSpot^TM^ 488 (Biotium, 70065), diluted 1:500 in live-cell imaging medium and incubated for 30 min. Images were acquired in the lipid (BDP or LipidSpot), tau_A647_ and, when applicable, Hoechst 33342 channels. Spot detection analysis was used to quantify the number of LDs. Where indicated, the number of lipid puncta was normalized to brain organoid area or to number of neuronal nuclei. Spot colocalization analysis was used to assess the number of BDP and tau-positive puncta.

#### Fluorescent lipid analogs

Prior to lipid loading, hiPSC-derived forebrain neuronal cultures were gently washed with serum-free media. Fluorescent lipid analogs were dissolved in the same serum-free media at a final concentration of 2.5 µM, except for BODIPY FL C12 CholEsteryl (Cholesterol ester, CE), which was used at 10 µM. BODIPY FL C12-Sphingomyelin (Sphingomyelin, SM) and TopFluor Cholesterol (Cholesterol, Chol) were applied for 1 h, whereas BODIPY 558/569 C12 (BDP-C12) and CE were incubated for 24 h in neurons preconditioned with 20 nM tau_A647_. Prior to imaging, neurons were rinsed once with serum-free media. Images were acquired for the respective fluorescent lipid analog, tau_A647_ and Hoechst 33342 channels. Colocalization between lipids and tau was quantified using Mander’s coefficient.

#### Modulation of neutral lipid storage

hiPSC-derived forebrain neuronal cultures were preincubated with DGAT inhibitors 1 (2.5 µM (Merck, A1737-1MG) and DGAT inhibitor 2, at 10 µM (Hölzel diagnostika, HY-110381- 5mg) for 4h in serum-free media. LD formation was then induced with oleic acid (OA; Merck, O1008-1G). OA was pre-complexed with BSA to reduce toxicity and improve delivery to neurons: 200 µM OA in 50% ethanol was combined with 10% fatty acid-free BSA (Merck, 820024-M) in PBS at a 6:1 molar ratio, together with 2 µM BODIPY 493/503, and incubated at 37°C for 1 h. The OA-BSA complex was subsequently diluted in serum-free medium and applied to cells for 24 h prior to live-cell imaging. Tau_A647_ (20 nM) was added concurrently with the OA–BSA mixture. Where indicated, DGAT inhibitors were maintained in the media throughout the experiment. A vehicle control containing an equivalent concentration of DMSO was included. Live-cell images were acquired in the BDP, tau_A647_ and Hoechst 33342 channels. LD (BDP puncta) and their colocalization with tau_A647_ were quantified using spot detection and colocalization analysis pipeline.

#### Tubulin imaging

hiPSC-derived forebrain neurons conditioned with 20 nM tau_A594_ were incubated with SiR700-tubulin (Spirochrome, SC014) at a final concentration of 1 µM for 1 h. Live-cell time-lapse imaging was performed in the tau_A594_ and tubulin channel to assess colocalization of tau trajectories with microtubules.

#### Dual-color tau puncta tracking

hiPSC-derived forebrain neuronal cultures or astrocyte cultures (hiPSC-derived astrocytes or CCF-STTG1 cells) were first labelled with CellTracker Green (Invitrogen, C2925) to asses weather tau is intracellular. CellTracker Green was incubated at 5µM final concentration for 15 min. Neurons were washed once and neuronal media was conditioned with 20 nM tau_A594_ for 2 h. Following a second rinse, neuronal media was conditioned with 20 nM tau_A647_ for 30 min. Cells were washed again prior to imaging. Time-lapse images were acquired in the CellTracker Green, tau_A594_ and tau_A647_ channels at 0.1Hz for 20 min. Colocalization of dual-color tau puncta was assessed at 20 min time point by spot detection and colocalization analysis.

#### Pharmacological dissipation of tau puncta

hiPSC-derived forebrain neurons conditioned 20 nM tau_A647_ were labelled with CellTracker Green (5µM, 15 min). After washing, the same field of view (FOV) was imaged prior and 10 min after addition of 1,6-hexaendiol (1,6-HD, Merck, 240117-50G) at a final concentration of 3% in imaging medium. Live-cell images were acquired in the CellTracker Green and tau_A647_ channels and tau puncta size was quantified following treatment.

#### TauSNAP imaging

O6-benzylguanine (BG) conjugated to either Abblive590 or Janelia Fluor 646 (JF646) were synthesized as previously described ^117^. To visualize tauSNAP subcellular localization in hiPSC-derived forebrain neurons from the tauSNAP PiggyBac hiPSC line, expression was induced by adding doxycycline 1 µg/ml final concentration (Thermo Fisher Scientific, J67043.AD)) to the neuronal maturation media 5 days prior to the experiment and maintained during routine media changes. On the day of imaging, neurons were incubated with either BG-Abblive590 or BG-JF_646_ at 1 µM in live-cell imaging media for 30 min, followed by three washes and a ≥4 h incubation in fresh imaging medium. Prior to live-cell imaging, co-staining with BDP or tubulin was performed as described above. Live-cell images were acquired in the tauSNAP, BDP or tubulin, and Hoechst 33342 channels. Colocalization between tauSNAP and BODIPY or tubulin signals was quantified using Mander’s coefficient.

#### Lipid peroxidation sensor

hiPSC-derived forebrain neuronal cultures were incubated with 2 µM BODIPY 581/591 C11 (BDP-C11) for 1h, followed by three washes prior to treatment with 20 nM tau_A647_. Where indicated, neurons were treated with NMDA (500µM, biotechne, 0114/500) or Erastin (50µM, Merck, 329600-5MG) for 4h. A vehicle control containing the same concentration of DMSO as used for the Erastin treatment was included. Prior to imaging, neurons were washed again with the same imaging medium. Images were acquired in the reduced (BDP-C11red), oxidized (BDP-C11ox), tau_A647_ and Hoechst 33342 channels. To compare the extend of colocalization between tau and BDP-C11ox or BDP-C11red, Mander’s coefficient was used, as the diffuse BDP-C11red signal is not suitable for spot detection analysis. The effect of NMDA treatment on the proportion of tau puncta colocalizing with BDP-C11ox puncta was further quantified using spot detection and colocalization analysis. Lipid peroxidation was quantified either as BDP- C11ox intensity normalized to neuronal nuclear count or as the ratio of BDP-C11ox/red.

#### For lipid peroxidation assay in brain organoid

brain organoids were incubated with 2 µM BDP-C11 for 24h in full maturation media. Lipid peroxidation was then induced by treatment NMDA (500µM) or Erastin (50µM) for 4h. A DMSO vehicle control was included. Organoids were subsequently washed three times, and the maturation media was conditioned with 20 nM tau_A647_ for 24h. Following a final wash, organoids were prepared for live-cell imaging. Images were acquired in the BDP-C11red, BDP-C11ox and tau channels, and lipid peroxidation was quantified as the ratio of BDP-C11ox/red.

#### Transfer of lipids from neurons to astrocytes

This protocol was adapted from previously published procedures (Jennifer Lippincott-Schwartz and colleagues^50^). hiPSC-derived forebrain neuronal cultures grown in 12 mm coverslips were incubated with BDP-C12 and, when indicated, 20 nM tau_A647_ in serum-free BrainPhys Image Optimized media for 24 h. The following day, neurons were washed twice and incubated in fresh imaging medium for 2 h to allow release of excess lipid. Co-culture between lipid-enriched neurons and untreated astrocytes was established by positioning the coverslip containing neurons face-down above astrocyte cultures (hiPSC-derived astrocytes or CCF-STTG1 cells) in 24-well plates. Cells were separated using paraffin wax spacers to prevent direct contact while allowing exchange of medium. Neurons and astrocytes were co-incubated for 4 or 24h. Co-cultures were then separated and imaged live to assess transfer of fluorescent lipids and tau from neurons to astrocytes. Images were acquired in the BDP-C12, tau_A647_ and Hoechst 33342 channels. BDP-C12 fluorescence intensity was quantified in both cell types and normalized to the cell number.

To asses transfer of peroxidized lipids, neurons were treated with BDP-C11. When applicable, neuronal media was first conditioned with 20 nM tau_A647_ for 24h. Neurons were then incubated with 2 µM BDP-C11 for 1h, washed three times, and treated with NMDA (500µM) or Erastin (50µM) for 4h. A DMSO vehicle control was included. Neurons were washed again and co-cultured with untreated astrocytes as described above for 4 or 24 h. Images were acquired in the BDP-C11red, BDP-C11ox, tau_A647_ and Hoechst 33342 channels. To quantify the extend of peroxidation after co-culture, we quantified BDP-C11ox intensity normalized to neuronal nuclear count or as the BDP-C11ox/red ratio in both neurons and astrocytes.

### Single-molecule measurements of lipids in neuronal media

Prior to lipid loading, hiPSC-derived forebrain neuronal cultures were washed once with serum-free live-cell imaging media. Cells were then incubated with 2.5 µM BDP-C12 together with 20 nM tau_A647_ for 24h. The following day, neurons were washed twice and incubated in fresh live-cell imaging medium for 2 h to allow release of excess lipid. The medium was subsequently replaced and cells were incubated for an additional 4 h to allow secretion of fluorescent lipid molecules. Where indicated, NMDA (500µM) was added to the live-cell imaging media. Single fluorescent lipid molecules in the medium were detected using fluorescence correlation spectroscopy (FCS). Measurements were performed on a MicroTime 200 time-resolved fluorescence system (PicoQuant) based on an inverted Olympus IX73 microscope (Olympus, Tokyo, Japan).

### Immunofluorescence staining

Brain organoids were fixed in PBS with 4% paraformaldehyde (PFA, Science Services, E15710), 1% glutaraldehyde (Merck, G5882-10×1ML) for 1 h at 4°C, followed by three washes in PBS (10 min each). Organoids were cryoprotected overnight in a 40% sucrose (Merck, S0389-1KG) in PBS. Following, organoids were embedded in a 10% sucrose/13 % gelatin (Merck, G2500-100G), and stored at −80 °C upon further use. Sections (14 µm) were cut using a CryoStar NX 70 cryostat (ThermoScientific), mounted on Superfrost Plus slides (Menzel, MZ-0031), and stored at −80 °C. Sections were washed with warm PBS, refixed for 10 min to dissolve and remove any remaining gelatin. Subsequently, sections were refixed (4% PFA, 0,1% glutaraldehyde) for 20 min at room temperature. Following three washes in PBS (10 min each), the sections were treated with 0.2% Triton-X100 (Merck, 93443) in PBS for 2.5 h. Following three washes with 0.05% Triton-X100 in PBS (10 min each), blocking was performed for 2 h with 10% normal donkey serum (abcam, ab7475) and 0.05% Triton-X100 in PBS. Primary antibodies were applied overnight at 4 °C in 5% normal goat serum with 0.05% Triton X-100 in PBS. The following primary antibodies were used: rabbit polyclonal anti-Sox2 (1:200 dilution, Merck, AB5603); mouse monoclonal anti-Tubulin ß3 (1:500 dilution, BioLegend, 801201) and rabbit monoclonal anti-S100 beta (1:100 dilution, Abcam, ab52642). Following 3x washes with 1% NGS in PBS-Triton-X100 for 30 mins, the sections were incubated with the appropriate secondary antibody in 5% NGS and 0.05% Triton-X100 in PBS for 2 h at 37°C. The following Alexa Fluor™ 594 -conjugated secondary antibodies were used (1:500 dilution): Goat anti-Rabbit IgG (H+L) Cross-Adsorbed Secondary Antibody (Thermo Fisher Scientific, A11012) and Goat anti-Mouse IgG (H+L) Cross-Adsorbed Secondary Antibody (Thermo Fisher Scientific, A11005). Following 3x washes with 1% NGS in PBS-Triton-X100 for 30 min, sections were washed 3x with PBS (5 min each). Nuclei were stained with DAPI (1µg/ml; Thermo Fisher Scientific, 62248) in PBS and incubated at room temperature for 15 min, followed by 3x washes in PBS. Sections were mounted using ProLong Gold Antifade Mountant (Thermo Fisher Scientific, P36934) and stored at 4°C.

For hiPSC-derived forebrain neurons, cells were fixed in PBS with 4% PFA, 0.1% glutaraldehyde for 10 min at room temperature. Subsequently, cells were washed 3x with PBS. Blocking was performed for 1 h with 2,5% BSA, 10% NGS (Biozol, ENG1000-100) or 10% NDS (Abcam, ab7475), depending on the host of the primary antibody, and 0.01% Triton-X100 in PBS. Primary antibodies were applied in blocking solution, and incubated overnight at 4°C. The following primary antibodies were used: rabbit polyclonal anti-PLIN2 antibody (1:200 dilution, Merck, HPA016607); rabbit monoclonal anti-Rab5 (1:200 dilution, Abcam, ab218624); mouse monoclonal anti-LAMP1 (1:200 dilution, DHSB, H4A3); goat polyclonal anti-APOE (1:250 dilution, Merck, 178479). Following 3x washes with PBS (10 min each), the cells were incubated with the appropriate secondary antibody in blocking solution for 1 h. The following secondary antibodies were used (1:1000 dilution): Donkey anti-Mouse IgG (H+L) Highly Cross-Adsorbed Secondary Antibody, Alexa Fluor™ 488, (Thermo Fisher Scientific, A21202); Donkey Anti-Rabbit IgG H&L (Alexa Fluor® 647) (Abcam, ab150075) and Donkey anti-Goat IgG (H+L) Highly Cross-Adsorbed Secondary Antibody, Alexa Fluor™ Plus 488 (Thermo Fisher Scientific, A32814). DAPI (1µg/ml) was included during secondary antibody dilution. Cells were washed 3x with PBS (10 min each) and mounted in IBIDI mounting media (ibidi, 50001).

### Thioflavin-S staining

Brain organoids media was conditioned with 20nM Tau594 or Tau fiber 594 for 24h. Next day, conditioned media was wash out and replaced for fresh media and 120h later organoids were fixed, embedded and cryosectioned following as described above. After permeabilization (0.25% Triton-X100, 30 min), sections were incubated for 15 min with freshly prepared Thioflavin-S (Biomol, ABD-23059) solution 0.05% Thioflavin-S w/s in 50% ethanol/water. The following series of washes (2x, 20 min with 50% ethanol, 1x, 20 min with 80% ethanol, 1x, 5 min with 0.05% TritionX100 in PBS and 1x, 5 min wash with water) were performed before mounting the sections in water and imaging directly.

### Fluorescence image acquisition

For all live-cell imaging experiments described below, images were acquired on a Dragonfly spinning disk confocal microscope (Andor, Oxford Instruments). Fixed samples were imaged using a Leica Stellaris 8- Falcon-STED confocal microscope (Leica Microsystems). FCS measurements were done on MicroTime 200 time-resolved fluorescence system (PicoQuant), which is also equipped with a two-photon laser (Spectra Physics) that was used for ThS imaging. FLIM measurements for EM correlation were performed on a Luminosa single photon counting (TCSPC) confocal fluorescence system (PicoQuant). Image acquisition parameters are described in detail in following sections:

#### Live-cell imaging

For all live-cell imaging experiments described above, images were taken using a Dragonfly spinning disk confocal microscope built on a DMi8 inverted microscope (Leica Microsystems) and equipped with an environmental (CO2 and temperature) control that is part of the Systems Biology Imaging platform at Max Delbrück Center (MDC) in Berlin. Image acquisition was performed using Fusion software (Andor, Oxford Instruments). Imaging data were acquired using a Leica HC PL APO 63×/1.30 NA glycerol immersion objective except for calcium imaging, which was performed with a Leica ACS APO 20×/0.60 NA multi-immersion using glycerol as immersion media. Excitation was provided by 405 nm, 488 nm, 561 nm and 643 nm laser lines. Optical sectioning was set using a 40µm pinhole disk configuration. Emission fluorescence selected by a quad-band dichroic beam splitter prism (405/488/561/647) and detected with a Sona-2.0B11 sCMOS camera (Andor, Oxford Instruments) after appropriate emission filtering including 450/50, 525/50, 600/50, 700/75 bandpass filters for each laser line. Image acquisition was controlled using Fusion software (Andor, Oxford Instruments). Images were acquired at 1300×1300 pixels with an effective pixel size of 0.162 µm/pixel and recorded at 16-bit depth. For each experiment replicate, laser power and camera exposure time were kept constant across conditions to allow for comparison of fluorescence signal. For tauA647 puncta single-particle tracking, images were acquired at 2 Hz for 1 min. Calcium oscillations were recorded at a frequency of 5 Hz for at least 3 mins.

#### Immunofluorescence stainings

Fluorescence images of fixed, immunostained samples, including brain organoid sections and hiPSC-derived forebrain neuron cultures, were acquired using a Leica Stellaris 8- Falcon-STED confocal microscope which is part of the Systems Biology Imaging platform at Max Delbrück Center (MDC). Image acquisition was controlled using a Leica LAS X software (Leica Microsystems). The system is equipped with a White Light Laser (WLL) and HyD detectors. Images were acquired using Leica HC PL APO 40×/1.30 NA oil immersion objective or Leica HC PL APO 40×/1.30 NA oil immersion objective. Excitation wavelength were selected using the tunable WLL depending on the fluorophore of interest and emission was collected using spectral detection settings optimized for each fluorophore combination to minimize spectral overlap. To further minimize spectral crosstalk between fluorophores, channels were acquired sequentially, starting from the longest to the shortest excitation wavelength. Images were collected at 12-bit depth, and the number of pixels was adjusted according to the zoom factor to achieve an optimized spatial resolution (0.059µm/pixel). When indicated, z-stacks were acquired with a step size of 0.5 µm. For each experimental replicate, acquisition parameters, including laser power and detector gain were kept constant across experimental groups to allow for comparison of fluorescence signal

#### FCS

FCS measurements were performed using a MicroTime 200 time-resolved fluorescence system based on an inverted Olympus IX73 microscope (Olympus). A 60x Plan-Apo/1.4-NA water-immersion objective and a 560 nm excitation laser were used. Acquisition was performed in pulsed mode at a repetition rate of 20MHz. Fluorescence emission was separated from laser excitation using a zt488/543/635/730rpc Dichroic mirror (Chroma), focused into the aperture of a 100-μm–diameter pinhole and directed into a single-photon SPAD detector equipped with HQ600/200M Band-Pass Filter (Chroma)). Fluorescence intensity fluctuations were recorded using SymphoTime software (PicoQuant) for 30 s (10 repetitions, 10 s intervals). Two independent FCS measurement were performed per condition at different FOVs.

#### FLIM

FLIM imaging was carried out on a PicoQuant Luminosa single photon counting (TCSPC) confocal fluorescence system). First, an overview of the neuronal cultures grown on EM grids covering several mm2 was acquired by performing a spiral scan in transmission mode. To obtain high resolution FLIM images, fixed neuronal samples were excited with 405, 560 and 640 nm lasers lines (PicoQuant) in pulsed mode at repetition rates of 20 MHz, 6.7 MHz sync rate, using the PIE mode. Emitted photons were collected with a 60x UPlanXApo/1.2-NA water-immersion objective, passed through a zt488/543/635/730rpc Dichroic mirror (Chroma) and was focused into a 67 µm diameter pinhole. Fluorescence emission was detected using single-photon counting SPAD detectors (Excelitas)) equipped with 445/30 ET, 600/50 ET, 690/70 H band pass filters (all from Chroma). Photon arrival times were recorded using MultiHarp160 TCSPC module. Images of 30 µm^2^ were acquired at 256 x 256 pixels with a dwell time of 30 µs. Frames were collected until 7000 photon count was reached. Data were analyzed by Luminosa Software for a fast quality control and via the NovaFLIM software (both from PicoQuant) for a more detailed analysis.

#### ThS imaging

ThS–stained samples were imaged using two-photon excitation microscopy. Two-photon imaging was performed using a tunable near-infrared laser (excitation wavelength 700 nm) to excite ThS, enabling deeper tissue penetration and reduced photobleaching. Emission signals were collected using 500/50 BP bandpass filters optimized for ThS fluorescence. Imaging parameters, were kept constant across conditions to allow for quantitative comparison of amyloid-associated fluorescence signals.

### Correlative fluorescence and scanning electron microscopy

Brain organoids were conditioned with 20 nM tauA647 in brain organoid maturation medium for 48h and fixed with 2% formaldehyde (Electron Microscopy Sciences, 15710) and 2.5% glutaraldehyde (Sigma-Aldrich, G5882-10) in 0.1 M phosphate buffer (pH 7.2) for 2 hours at room temperature. Samples were embedded in low-melting agarose (Merck, A9414-10G) and sectioned (200 µm) using a vibratome. Sections were high-pressure frozen (HPF) using the Leica EM ICE (Leica Microsystems) using 3 mm and 6 mm wide, 300 µm deep gold-coated aluminium carriers (Wohlwend GmbH). To prevent ice crystal formation during HPF, 20% Ficoll PM 70 (Sigma-Aldrich, F2878-50) in 0.1 M HEPES pH 7.2 was used as a cryo-protectant. Freeze substitution was carried out using the Leica AFS2 with the freeze substitution processing unit (FSP). Resin fluorescence was preserved by freeze substitution with 0.1% uranyl acetate in glass distilled dry acetone and embedding in Lowicryl HM 20 resin (Polysciences) as previously described by (Ronchi et. al, 2021). The detailed workflow for the Leica-FSP run is described in Fig. SI4f. Resin blocks were sectioned (80 nm) using a Reichert Ultracut S microtome and collected on glass coverslips, which were carbon coated using the Safematic CCU-010 carbon coater (BALTIC Praeparation e.K.). An alphabetical grid mask (Leica Microsystems) was used to create a system of coordinates to facilitate correlation between imaging modalities. Fluorescence intensity was acquired using the Luminosa microscope (PicoQuant GmbH). Subsequently, sections were post-stained with 2% aqueous uranyl acetate and 3% Reynold’s lead citrate and imaged using the Helios 5CX HydraDualBeam system (Thermo Fisher Scientific). Data acquisition and correlative analysis of fluorescence and scanning electron microscopy (SEM) data was performed using Maps software version 3.27 (Thermo Fisher Scientific). Tile scans were collected using an acceleration voltage of 2kV and a beam current of 0.4-0.8 nA. Overview tile scans of the alphabetical grid and sections were aquired using secondary electrons detected using an Everhart–Thornley detector (ETD). Higher magnification tile scans of the tissue were collected using backscattered electrons detected by a CBS detector. Pixel dwell times of 1-3 µs were used with image resolutions of 3072×2048 or 6144×4376 pixels.

### Correlative fluorescence lifetime and transmission electron microscopy

hiPSC-derived forebrain neurons were cultured on PLO/laminin-coated 35 mm glass-bottom dishes with an imprinted 50 µm coordinate grid (ibidi, 81148). Neurons were incubated with BODIPY-C12 in serum-free BrainPhys medium for 24 h. Following a wash with warm PBS to remove excess non-internalized lipids, neurons were conditioned with 20 nM tauA647 for another 24 h. Neurons were then washed again in warm PBS to remove non-internalized excess tau protein and returned to full maturation media for 24 h prior to fixation. Cells were fixed with 4% formaldehyde, 0.5% glutaraldehyde in 0.1 M phosphate buffer (pH 7.2). Fluorescence Lifetime Imaging (FLIM) was performed using the Luminosa microscope (PicoQuant GmbH). Following FLIM, samples were processed for electron microscopy. Cells were incubated on ice for 1 hour in 1.5% potassium hexacyanoferrate (II) trihydrate (K_4_Fe(CN)_6_ · 3H_2_O) (Merck, 9387) and 2% osmium tetroxide (OsO_4_) (Merck, 75632) in 0.1 M phosphate buffer (pH 7.2). After washing with buffer, the cells were incubated for 2 min in 0.1% aqueous thiocarbohydrazide (TCH) (Merck, 223220) and then rinsed with MilliQ water. A second osmium treatment (1 % aqueous OsO_4_) was performed for 30 min on ice. Following another MilliQ water rinse, samples were stained with 1% aqueous uranyl acetate for 1 h at 4°C. Dehydration was carried out through a graded acetone series (30%, 50%, 70%, 90% and twice in 100% acetone). The cells were then gradually infiltrated with epoxy resin (Polysciences, Polybed812, 08791) using 30% and 70% epoxy resin–acetonel mixtures, followed by an overnight incubation in 100% resin. Subsequently, coverslips were detached from the plastic dishes and positioned, cell-side down, onto BEEM specimen embedding capsules (Ted Pella, 133-P) pre-filled with resin. Samples were polymerized for 48 h at 60°C. Finally, the glass was removed from the resin by using liquid nitrogen, and the resulting blocks were sectioned with a Reichert Ultracut S microtome. Sections (180–200 nm) were collected on carbon-coated Formvar slot grids and imaged using a JEOL 2100 Plus TEM (JEOL, Germany) operating at 200 kV acceleration voltage, convergence angle α3, spot size 1-2. 2D electron micrographs were recorded using the XAROSA CMOS camera and RADIUS 2.2 software (EMSIS, GmbH). Tilt series were acquired from - 60° to +60° at 1° tilt increments using SerialEM version 4.1.14^118^, Fiducialess alignment and tomogram reconstructions were done using weighted backprojection in IMOD version 5.1.0^119^.

### Imaging data analysis

Image processing and analysis was performed primarily using ImageJ/Fiji^120^ unless otherwise stated. As standard preprocessing step, background correction was applied. Background correction was applied by subtracting measured background fluorescence. Quantitative measurements were normalized to the nuclear count or organoid area where indicated. Nuclei counts and area measurements were obtained using the “Analyze Particle” function in ImageJ/Fiji. Nuclei were counted based on the Hoechst or DAPI signal and size (1,000-10,000) and circularity (0.5-1.0) thresholds. Organoid area was determined from BODIPY 493/503 signal using non-restrictive thresholding to capture the full signal-positive area.

#### Radial quantification of tau spreading in brain organoids

To analyze the translocation of tau_A647_ signal in brain organoids upon spreading, image analysis was performed on confocal images of brain organoid sections. Raw image data were processed using Fiji (ImageJ). For each time point, organoids were segmented based on fluorescence signal to define the outer boundary. The radial intensity profiles of tau_A647_ fluorescence were calculated from the organoid surface toward the core. Line projections (30 of them) were drawn from the organoid surface to the core. To quantify tau spreading, line profiles were defined at increasing depths from the surface. Intensity distributions were compared across time points (0-120 h) to assess spatial redistribution of tau_A647_. The progressive inward translocation was quantified by shifts in peak fluorescence intensity from peripheral to central regions. The depth of tau_A647_ spreading was determined as the maximum distance from the organoid surface at which fluorescence intensity exceeded background levels. Cell layer equivalents were estimated by the average number of nuclei crossed by the line profile, determined from the nuclear staining. All measurements were performed on at least three independent organoids per condition. Data are presented as mean ± standard deviation.

#### Calcium imaging

To identify calcium activity sources, temporal maximum projections were segmented using a Cellpose model that we trained on our 3D organoid data (Cellpose version: 4.0.8; platform: darwin; python version: 3.12.10; torch version: 2.10.0). The resulting segmentation mask was filtered to remove objects deemed too small (minimum size: 200 pixels), too big (maximum size: 2000 pixels) or too eccentric (maximum eccentricity: 0.98). For each segmented object, the fluorescence time trace (F) was obtained by averaging pixel intensities across all the objects’ pixels at each time point. The raw time trace F was then normalized to the baseline (F_0_), estimated as the 10th percentile within a 10 s rolling window, to compute ΔF/F_0_=(F-F_0_)/F_0_. Peaks in the normalized traces were identified using the ‘find_peaks’ function (height threshold: 0.025; prominence threshold: 0.025) from the ‘SciPy’ library (version 1.17.0). The average firing frequency per trace was calculated as the number of peaks divided by the total recording duration.

#### Intensity-based colocalization

Colocalization analysis was performed in ImageJ/Fiji using JaCOP plugin^121^. Brightness and contrast were adjusted automatically and consistently across all images to facilitate consistent comparison between experimental replicates. Thresholds for tau_A647_ channel (M1) were determined automatically. For the second channel (M2, lipid or subcellular marker signal), fixed thresholds were applied uniformly across all images. Mander’s colocalization coefficient (M1 and M2) were calculated to quantify the fraction of tau channel (M1) overlapping the second channel (M2). Coefficients were calculated per image and averaged across conditions.

#### Fluorescence intensity

Fluorescence intensity was quantified on ImageJ/Fiji using the “Measure” function on background-subtracted images (no brightness/contrast adjustments applied). Mean gray values were used as a readout. When indicated, fluorescence intensity was normalized to organoid area or number of neuronal or astrocytic nuclei (cell number). For analysis of lipid transfer between neurons and astrocytes, fluorescence intensity was normalized using min-mas scaling for each experimental replicate to account for inter-experimental variability. The BDP-C11 oxidation ratio was calculated as the ratio of BDP-C11ox to BDP-C11red intensity.

#### Spot detection and coordinate-based colocalization

In ImageJ/Fiji, brightness/contrast were adjusted automatically and uniformly across all images to ensure consistent comparison between experimental replicates. For modulation of neutral lipid storage experiments, brightness/contrast of the BDP channel was defined based on the parameters determined for OA treatment condition and applied to the rest of the conditions. Tau_A647_ puncta were then detected using the RS-FISH plugin^122^ in ImageJ/Fiji, employing a radial symmetry-based spot detection method, with the following parameters: sigma = 2, threshold = 0.01 support region radius = 4, inlier ratio = 0.1, maximum error = 1.5, and intensity threshold = 20000. Neutral lipid puncta, identified based on BODIPY 493/503 or BODIPY 581/591 C11 green fluorescence signal, were detected using the same approach and same parameters (threshold =0.035). Centroid XY coordinates were extracted for both spot populations. The mean number of puncta per image was normalized by 100 µm^2^ of organoid cross-sectional area or by number nuclei in neuronal cultures.

Coordinate-based colocalization between tau_A647_ and neutral lipid puncta was performed using a custom Python package (fish-colocalization, https://gitlab.com/ida-mdc/fish-colocalization). The two sets of spot coordinates per image were matched using a greedy nearest-neighbor algorithm (scipy.spatial.KDTree)^123^: iteratively, the closest pair of unmatched spots across the two channels was assigned as colocalizing, provided their Euclidean distance did not exceed 0.65 µm, in accordance to half of the diameter of neutral lipid puncta, which was around 1 µm. Each spot could be matched at most once. To calculate colocalization between tau_A647_ and PLIN2 puncta, we adjusted the maximum distance allowed between two colocalizing puncta to a shorter distance (0.235 µm) as the diameter of PLIN2 puncta didn’t exceed 500 nm. The percentage of tau_A647_ puncta with lipid puncta (BODIPY 493/503 or BODIPY 581/591 C11 green) or the percentage of lipid puncta colocalizing with tau_A647_ puncta was calculated relative to the total number of detected tau_A647_ puncta or BODIPY 493/503/BODIPY 581/591 C11 green, respectively. The percentage was calculated per each image and averaged across conditions. Coordinate-based colocalization between tau_A647_ and tau_A488_ in the dual-color tau puncta tracking was detected, measured and calculated following the same pipeline.

#### Single-particle tracking

Trajectory analysis of tau puncta was performed using TrackMate^124^ (ImageJ/Fiji, v8.1.6). Time lapse images were processed in ImageJ/Fiji to correct for bleaching with the “Bleach Correction” function and the “Simple Ratio” correction method. Brightness/contrast were adjusted automatically and equally in all the images in ImageJ/Fiji to facilitate consistent comparison between experimental replicates. Tau_A647_ puncta were segmented using the Stardist detector^125^ with default parameters with an estimated radius of 0.82 µm and a maximum intensity of 36,000 a.u. Trajectories were reconstructed using a LAP (Linear Assignment Problem mathematical framework) tracker with maximum linking distance of 3.5 µm. Gap closing up to 4 frames was allowed with 4.5 µm as a maximum distance. Tracks shorter than 20 frames were excluded from the analysis. For each trajectory, instantaneous speed was calculated as the displacement between consecutive frames divided by the time interval. The maximum instantaneous speed per track and the mean velocity, as the total displacement from the first to the last frame, were calculated.

#### Colocalization with microtubules

In ImageJ/Fiji, the average intensity of the time-frame images of the tubulin channel were projected in Z. Brightness/contrast were adjusted automatically and uniformly across all images to ensure consistent comparison between experimental replicates and a binary mask was generated using the Huang automatic thresholding algorithm. Tau_A647_ trajectories were calculated using TrackMate following the previously described pipeline and the detected spots corresponding to the 100 tracks with higher displacement values were exported as ROIs for ImageJ/Fiji. ROIs were loaded into the tubulin binary mask and their corresponding mode value was measured. The percentage of tau_A647_ puncta overlapping with microtubules was calculated by counting the number of spots for which the mode value was 255 (=overlapping tubulin signal) in respect to the total amount of spots detect for each individual track.

#### FLIM

FLIM measurements were analyzed using SymPhoTime or NovaFLIM softwares, both from PicoQuant. Photon arrival time histograms were generated for each pixel and decay curves were fitted using a bi-exponential decay model to account for heterogeneous fluorophore environments. The instrument response function (IRF) was calculated and included in the fitting procedure. Fits were performed using a least-squares reconvolution method. From the fitted decay curves, the short (τ_₁_) and long (τ_₂_) lifetime components, as well as their respective amplitude fractions (A_₁_ and A_₂_), were extracted. The amplitude-weighted mean lifetime (τ_mean) was calculated as:

τ_mean = (A_1_T_1_ + A_2_τ_2_)/(A_1_ + A_2_)

Lifetime maps were generated for visualization, applying consistent color scales across all experimental conditions. Where indicated, regions of interest (ROIs) were manually defined or based on segmentation masks, and mean lifetime values were extracted for quantitative comparison.

#### Tau puncta and lysosome size measurements

Tau_A647_ or Lamp1-positive puncta segmentation was performed using StarDist plugin in ImageJ/Fiji (default detection parameters). As preprocessing step, brightness/contrast were adjusted in all channels for consistent comparison between experimental replicates. Object size was defined as the segmented area. For tau_A647_ puncta analysis, objects with an area > 1.5 µm^2^ were not included. For lysosomal size analysis, segmented objects with mean fluorescence intensity < 25,000 a.u. were excluded. Mean puncta size was calculated per image and averaged across conditions.

#### FCS analysis (number of molecules and diffusion time)

FCS intensity traces were used to calculate autocorrelation curves. For each measurement, autocorrelation curves were calculated (10 × 30 s per measurement) and averaged together to obtain statistical variations. The average autocorrelation curves were fitted to a 3D Brownian diffusion model, volume weighted by the inverse square of the SD:

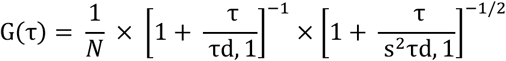

where G(τ) is the autocorrelation function for translational diffusion as a function of time τ, N is the average number of fluorescent molecules in the laser focal volume, and τD is the translational diffusion time of particles. To quantify lipid secretion, both the diffusion time, τD (Fig. SI5c), and the number of molecules, N (Fig. 5g) of the detected BDP-C12 fluorescence were analyzed.

#### Morphometry of tau inclusions

Scanning and transmission electron micrographs were analyzed to quantify the circularity and area of tau-positive inclusions using Fiji (NIH). A Wacom Intuos S graphics tablet was used for manual tracing of correlated tau inclusions, and measurements were performed in Fiji.

### Statistics

Statistical tests were conducted in GraphPad Prism 9 for Windows 9.4.1. Data are represented as mean ± SD and visualized as bar plots with individual data points overlaid, unless otherwise indicated. Ca imaging data (Fig. SI1 a-e) and FCS measurements (Fig. 5g) are presented as median and interquartile ranges in violin plots. Each dots represents sample number (*n*); in Figure 1 and 2, biological replicates and in Figures 3 to 6, technical replicates. No statistical methods were used to predetermine sample size, but sample sizes are consistent with those commonly used in the field. When appropriate, data were normalized within each experimental replicate using min-max scaling to account for inter-experimental variability across independent experiments. Data distribution was evaluated by visual inspection of Q-Q plots combined with normality testing, selected according the sample size (Shapiro-Wilk test for N > 10 and D’Agostino-Pearson for N < 10). Homoscedasticity of variances was evaluated using an F-test. When data met normality assumptions and equal distribution, two-tail t-test was used to compare two groups (paired for related samples or unpaired for independent groups), one-way ANOVA to compare means across > 3 groups or two-way ANOVA to compare the influence of 2 different factors (followed by a Tukey’s post-hoc test). Data showing unequal variances was evaluated by Welch’s test. For non-normally distributed data, non-parametric tests were used: the Mann–Whitney U test for two unpaired group comparisons, the Wilcoxon test for two paired group comparisons and the Kruskal–Wallis test followed by Dunn’s multiple comparisons test for more than two groups. All tests were two-sided, and a significance threshold of *p* < 0.05 was applied.

Exact statistical tests, sample sizes (*n*), and definitions of biological and technical replicates are specified in the corresponding figure legends. Statistical significance is indicated as follows: *p* < 0.05 (*), *p* < 0.01 (**), *p* < 0.001 (**), and *p* < 0.0001 (****). For clarity, only statistically significant differences are annotated in the figures.

## Author contribution

M.B and A.O.M conceptualized the study and designed the experiments, A.O.M. performed, analyzed and curated the data for most the experiments of the study, supervised by M.B. and with contributions from: D.I.; performed part of the flow cytometry experiments and analysis of all flow cytometry data, M.C.N.; generated Tau-SNAP line, G.M; obtained lipidomics data, M.F and L. M. S. P. develop the code for lipidomics analysis and perform the analysis with support and supervision by S.K. M.A and K.H analyzed and curated transcriptomics data, M.C.R performed the EM part of the FLIM-EM pipeline, analysis and data curation supervised by S.K. L.D.J develop the code for Ca imaging data analysis, analyzed and curated Ca imaging data with the support of M.P and supervised by J.M. J.R purified and labeled the tau protein, performed immunofluorescence staining, viability assays and provided support for cell culture. M.B performed ThS experiments and analysis, radial tau spreading analysis and curated FLIM data from the FLIM-EM pipeline. C.C.J. and N.R. provided primary mouse neuronal cultures. E.B developed the code for spot colocalization with supervision by D.S. J.B provided SNAP-tag ligands. A.R.W provided forebrain organoids, advice on transcriptomics experiments and performed RNA purification. M.B. provided funding acquisition. A.O.M and M.B. wrote the original draft and all authors participated in the review process.

## Acknowledgements

We thank all members of the Birol laboratory for their support, insightful discussions, feedback, and continuous encouragement throughout this project. We are especially grateful to Elisabeth Fritsch and Paula Santos-Otte for sharing automation scripts and computational support. We thank Suzanne Wegmann, Thomas E. Willnow, Erich Wanker, Elizabeth Rhoades for valuable feedback, discussions, and advice during the development of this work. We are grateful to Sebastian Diecke and Silke Frahm for providing iPSC cell lines and guidance on stem cell culture. We also thank the Siffrin laboratory for sharing astrocyte differentiation protocols. We acknowledge the support of the imaging platform for microscopy assistance and technical expertise. We thank Elizabeth Rhoades for sharing tau plasmids and Ivano Legnini for PiggyBac and transposase plasmids. We thank Caroline Braeuning and the Genomics Technology Platform for assistance and support with FACS experiments and analysis and RNA sequencing. We additionally thank PicoQuant, Evangelos Sisamakis, Olaf Schulz, and Isabel Gross for discussions, support, and feedback related to the FLIM-EM. This project has received funding from the European Union’s Horizon Europe Framework Programme (deuterON, grant agreement no. 101042046 to JB). This work was supported by the Alzheimer Forschung Initiative (23047R) and the Alzheimer’s Association (AARG-22- 971947) through funding awarded to Melissa Birol.

